# Conservation of C4BP-binding Sequence Patterns in *Streptococcus pyogenes* M and Enn Proteins

**DOI:** 10.1101/2024.04.22.590534

**Authors:** Piotr Kolesiński, Matthew McGowan, Anne Botteaux, Pierre R. Smeesters, Partho Ghosh

## Abstract

Antigenically sequence variable M proteins of the major bacterial pathogen *Streptococcus pyogenes* (Strep A) are responsible for recruiting human C4b-binding protein (C4BP) to the bacterial surface, which enables Strep A to evade destruction by the immune system. The most sequence divergent portion of M proteins, the hypervariable region (HVR), is responsible for binding C4BP. Structural evidence points to the conservation of two C4BP-binding sequence patterns (M2 and M22) in the HVR of numerous M proteins, with this conservation applicable to vaccine immunogen design. These two patterns, however, only partially explain C4BP-binding by Strep A. Here, we identified several M proteins that lack these patterns but still bind C4BP, and determined the structures of two, M68 and M87 HVRs, in complex with a C4BP fragment. Mutagenesis of these M proteins led to identification of amino acids that are crucial for C4BP-binding, enabling formulation of new C4BP-binding patterns. Mutagenesis was also carried out on M2 and M22 proteins to refine or generate experimentally grounded C4BP-binding patterns. The M22 pattern was the most populated among M proteins, followed by the M87 and M2 patterns, while the M68 pattern was rare. These patterns, except for M68, were also evident in numerous M-like Enn proteins. Binding of C4BP via these patterns to Enn proteins was verified. We conclude that C4BP-binding patterns occur frequently in Strep A strains of differing M types, being present in their M or Enn proteins, or frequently both, providing further impetus for their use as vaccine immunogens.

## Introduction

*Streptococcus pyogenes* (group A *Streptococcus* or Strep A) is a widespread gram-positive bacterial pathogen that is responsible for morbidity and mortality on a global scale (1). While infection by Strep A usually results in mild conditions (e.g., pharyngitis or impetigo), a significant number of cases progress to acute invasive disease, such as necrotizing fasciitis or streptococcal toxic shock syndrome, which have high mortality rates (2). In addition, prolonged Strep A infection can lead to severe autoimmune sequelae (e.g., acute rheumatic fever) (3, 4), which continues to afflict populations in developing nations (5).

Strep A is highly proficient at avoiding phagocytic killing by the immune system. A frequent means by which Strep A achieves this is by recruiting soluble human C4b-binding protein (C4BP, ∼570 kDa) to the bacterial surface (6–8). Recruitment of C4BP inhibits deposition of the complement system opsonin C3b (by the classical and lectin pathways) on the bacterial surface, thereby antagonizing clearance of Strep A by phagocytic immune cells. A primary recruiter of C4BP is the essential virulence factor M protein (9, 10), which is antigenically sequence variable, with >220 M types having been identified (11). Despite this variability, C4BP-binding is pervasive in Strep A, with one study identifying C4BP-binding in 90 of 100 strains of differing M types (9).

M proteins form parallel, dimeric α-helical coiled coils which are attached to the bacterial cell wall and extend ∼500 Å into the extracellular space (12). The N-terminal 50 amino acids of mature M proteins (i.e., with the signal sequence cleaved) extend furthest into the extracellular space and are especially variable (called the hypervariable region, HVR). The HVR defines the M type. Surprisingly, the HVRs of several M types, despite lacking sequence conservation, are responsible for binding C4BP (6, 9, 13, 14). One mode of interaction between sequence-divergent M protein HVRs and C4BP was revealed by X-ray crystallography (10). Structures of four different M protein HVRs (M2, M22, M28, and M49) bound to a C4BP fragment (composed of the α1 and α2 domains, C4BPα1-2) revealed a common three-dimensional (3D) arrangement of M protein amino acids that interact with a common set of C4BP amino acids. The C4BPα2 domain supplies four sites forming a quadrilateral composed of a (1) hydrophobic pocket (denoted ΦHis 67) that binds large hydrophobic M protein amino acids; (2) the main chain amide of H67 (denoted N-His 67) that forms a hydrogen-bond with an M protein amino acid; and (3) R64 and (4) R66 (Fig. 1A). Arginines are versatile in providing a polar contact through their guanidinium group or hydrophobic interaction through their alkyl carbons. C4BPα2 R64 and R66 form polar contacts with M protein amino acids (D, E, or N) in M2, M22, and M28 HVRs, but in the M49 HVR, R64 forms hydrophobic contacts with hydrophobic M49 amino acids. The C4BPα1 domain provides a single site, the Arg39 nook, which consists of (5) a hydrophobic pocket formed by the main and side chain of R39, which interacts with a large hydrophobic M protein amino acid. In the case of the M2 and M49 HVRs, R39 also forms (6) salt bridges with negatively charged M protein amino acids.

**Figure 1.**
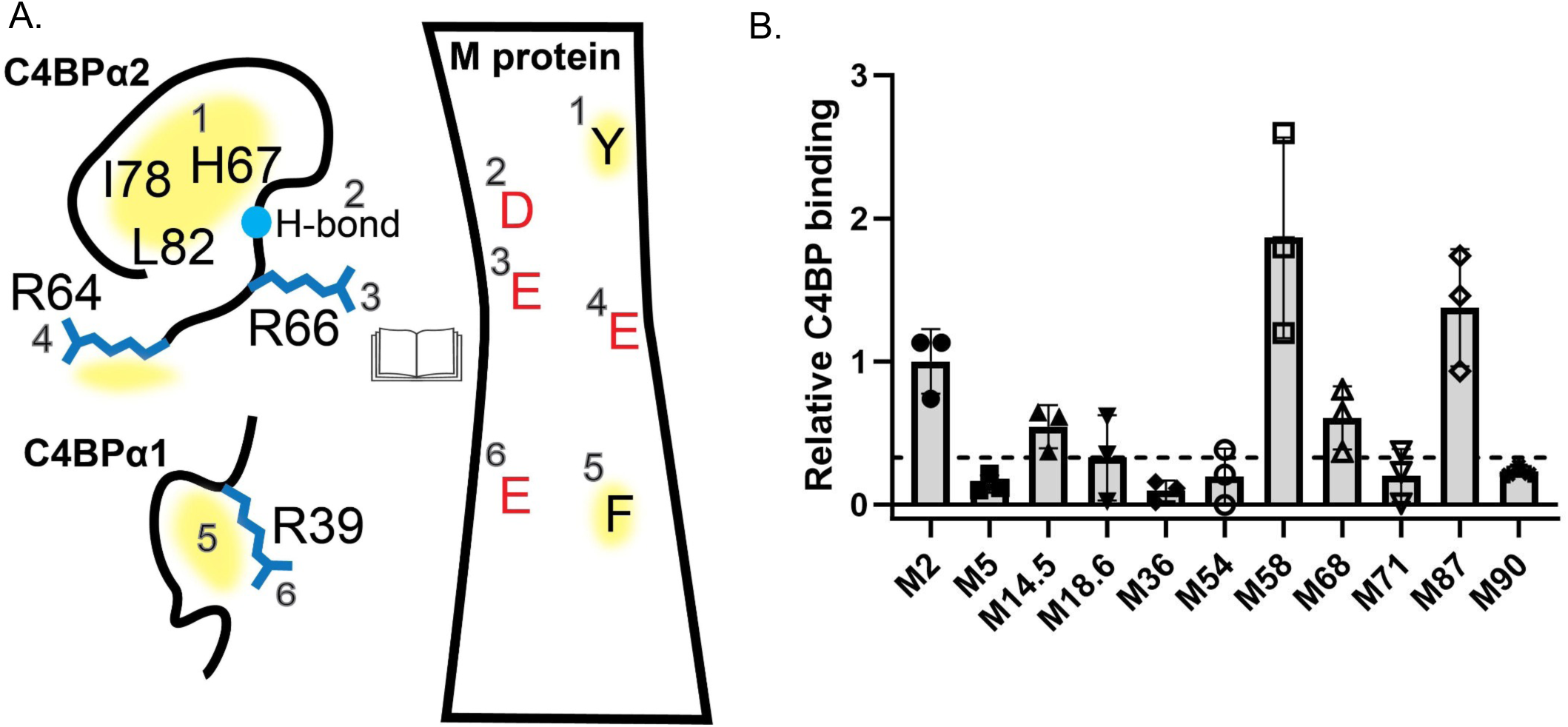
Binding of C4BP to M proteins. **A.** Schematic of C4BP sites (left) and M protein amino acids (right) involved in interaction, in open book format. The M protein amino acids do not correspond to any specific M protein but show interactions common to M2, M22, M28, and M49 HVRs. Yellow shading corresponds to hydrophobic interactions. **B.** Binding of soluble His_6_-M^N100^ protein constructs to immobilized intact C4BP, as evaluated by ELISA. Bound His_6_-M^N100^ protein constructs were detected with HRP-conjugated anti-His_6_ antibodies. Values were normalized by the value of C4BP-binding to His_6_-M2^N100^. Means and standard deviations from three biological replicates carried out in triplicate are shown. The dotted line shows the normalized value for C4BP-binding to His_6_-M5^N100^ plus three standard deviations.

While the C4BP-interacting amino acids in all four M protein HVRs are positioned similarly in three-dimensions, they differ in their heptad positions. Coiled-coil proteins have diagnostic heptads (*abcdefg*) in their primary sequence, in which the *a* and *d* positions are typically small hydrophobic amino acids that form the hydrophobic core of the structure. The heptad pattern of C4BP-binding amino acids is similar for the M2 and M49 HVRs and defines the M2 pattern for C4BP-binding, while the heptad pattern is similar for the M22 and M28 HVRs and defines the M22 pattern. The M2 and M22 patterns are evident in numerous M proteins, providing an explanation for how many Strep A strains bind C4BP (9, 10). The conserved M2 and M22 patterns occur in the midst of hypervariability and were thus unrecognized prior to structural studies.

Conservation of the C4BP-binding 3D pattern suggested its utility as a vaccine immunogen. Currently there is no vaccine against Strep A (15), with the sequence variability of M proteins being one of the major impediments (16–20). We found that an immunogen consisting of the C4BP-binding 3D pattern of M2 protein is capable of eliciting antibodies that react with M2 protein and are cross-reactive against other M protein types sharing the 3D pattern but not against those lacking it (21). These cross-reactive antibodies promote the opsonophagocytic killing of Strep A strains (21).

The M2 and M22 patterns only partially explain C4BP-binding. More than half of the 90 Strep A strains shown to bind C4BP lack an M2 or M22 pattern (9, 10). We therefore asked whether additional C4BP-binding patterns exist in M proteins. We identified C4BP-binding in several M proteins that lack an M2 or M22 pattern, and determined the structures of two, M68 and M87 protein HVRs, in complex with C4BPα1-2. Based on mutagenesis of these M proteins and effects on C4BP-binding, we constructed M87 and M68 C4BP-binding patterns. Similar mutagenesis and C4BP-binding experiments were carried out for M2 and M22 proteins, enabling refinement or generation of experimentally based M2 and M22 patterns, respectively. The M2, M22, and M87 patterns were also identified and verified in numerous M-like Enn proteins. Our results indicate that C4BP-binding patterns are highly prevalent in M and Enn proteins.

## Results

### C4BP-binding

We tested nine M proteins that lack an M2 or M22 pattern for C4BP-binding; these were all from strains shown to bind C4BP (9, 10). The M proteins were M14.5, M18.6, M36, M54, M58, M68, M71, M87, and M90. His_6_-tagged constructs consisting of the N-terminal 100 amino acids of these M proteins (designated by superscripted ‘N100’) were expressed in *E. coli* and purified, and assayed for binding to intact C4BP by ELISA (Fig. 1B). The binding of His_6_-M2^N100^ to C4BP was used for normalization. His_6_-M5^N100^ was used as a negative control, as M5 protein does not bind C4BP (21). We distinguished binders from non-binders using a threshold of three standard deviations above the C4BP-binding of His_6_-M5^N100^. His_6_-tagged M14.5^N100^, M58^N100^, M68^N100^, and M87^N100^ crossed this threshold, while the other five of the 9 did not (Fig. 1B). Co-crystallization with C4BPα1-2 was pursued with the four C4BP-binding M^N100^ protein constructs, and M68^N100^ and M87^N100^ yielded co-crystals that diffracted sufficiently for atomic-level modeling of the binding interface.

### M87–C4BP Interactions

The structure of the complex between M87^N100^ and C4BPα1-2 was determined by molecular replacement to 2.7 Å resolution limit (Supp. Table S1). The structure revealed the same global architecture observed previously (10), with two C4BPα1-2 molecules bound by dimeric, α-helical M87^N100^ (Fig. 2A). C4BPα1-2 was unchanged in conformation (rmsd 0.74 - 1.18 Å) (10), although certain loops distant from the interface with M87 were flexible and not modeled. The two M87 helices (amino acids, aa, 46-110 and 46-125) were asymmetric, with one less coiled than the other. Nevertheless, the ∼2.5 heptads of M87 protein from D80 to D96 that bound C4BP were closely superimposable (0.68 Å rmsd) (Supp. Fig. S1). This section was stabilized by knobs-into-hole packing, except at Y81 (*a* position), which faced outward from the coiled-coil core and contacted C4BP (Figs. 2B and C). The two M87-C4BP interfaces were similar (0.57 Å rmsd) but not identical. The buried surface area for one interface was 1498 Å^2^ and 956 Å^2^ for the other. Closer inspection revealed that the higher value was due to amino acids that were not in contact (>4 Å apart) but nevertheless excluded water at their interface.

**Figure 2.**
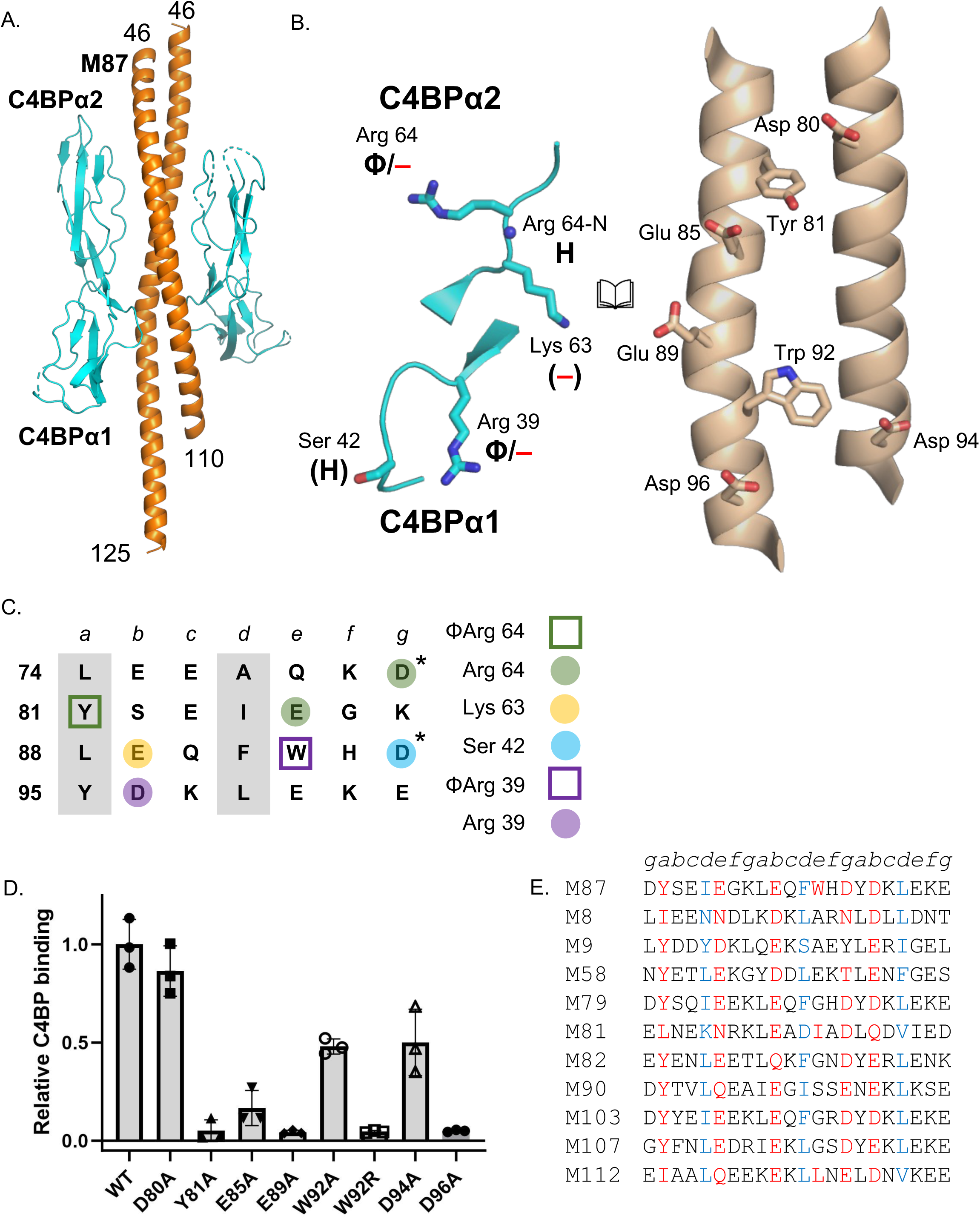
M87-C4BP interactions. **A.** Structure of M87^N100^ (orange) in complex with C4BPα1-2 (cyan), in cartoon representation. Loops of C4BPα1-2 that were not modeled are shown as dotted lines. **B.** Interaction between C4BPα1-2 (left) and M87 (right), in cartoon (main chain) or bonds (side chains) representation, shown in open book format. Carbons are cyan for C4BPα1-2 and wheat for M87; for both, nitrogens are blue, and oxygens red. Chemical character of interactions denoted as follows: Φ, hydrophobic; —, negative; H, hydrogen bond. Contacts that exceeded hydrogen-bonding or salt-bridging limits (≤3.5 and ≤4.0 Å, respectively) have these labels in parentheses. **C.** Heptad register of M87 74-101 (*a* and *d* positions in gray columns). Amino acids marked with an asterisk belong to the minor helix. Shapes indicate type of contact (square for hydrophobic, Φ; circle for polar), and coloring indicates which C4BP amino acid (depicted in legend to the right) was contacted. **D.** Binding of soluble, intact His_6_-M87 protein constructs to immobilized intact C4BP evaluated by ELISA. Bound His_6_-M87 protein was detected with HRP-conjugated anti-His_6_ antibodies. Values were normalized by the value of C4BP-binding to wild-type His_6_-M87. Means and standard deviations from three biological replicates carried out in triplicate are shown. **E.** Partial list of sequences of M proteins with M87 C4BP-binding patterns. Heptad positions are indicated above the sequences. Red indicates an observed (M87) or potential (other M proteins) C4BP-binding amino acid, and blue a *d* heptad position. A full list is in Supp. Table S2.

The buried surface areas considering only the amino acids that were in contact distance (≤4 Å) were more similar, 583 and 600 Å^2^. The surface complementarities of the two interfaces were 0.61 and 0.69, similar to values observed for other M protein HVR-C4BP complexes (10).

The interface was dominated by negatively charged M protein amino acids interacting with positively charged C4BP ones, supplemented with a few hydrophobic contacts (Figs. 2B and C). Each C4BPα1-2 molecule contacted predominantly one of the two M87 helices, which we termed the ‘major’ helix and the other the ‘minor’ helix. The C4BPα2 quadrilateral (Fig. 1A) as a whole was not involved in interaction, except for R64, which formed hydrophobic contacts using its aliphatic chain with M87 Y81 of the major helix and a salt bridge with D80 of the minor helix; Y81 was the only core *a* or *d* position amino acid involved in contacts. C4BPα2 R64 was also proximal to M87 E85 of the major helix, and with slight rearrangement would be within salt bridging distance (≤4 Å) of this amino acid. In one of the interfaces, the main chain amide of C4BPα2 R64 formed a hydrogen bond with M87 E85 of the major helix. C4BPα2 K63, which was not observed previously to have a role in M protein interaction (10), was within near-salt bridging distance of M87 E89 of the major helix. In contrast to the C4BPα2 quadrilateral, the C4BPα1 Arg39 nook had the same role as previously observed in the M2 pattern (10), forming hydrophobic contacts and a salt bridge to M protein. Here the former was to M87 W92 of the major helix and the latter to D96 of the minor helix. In one of the interfaces, the main chain carbonyl of C4BPα1 R39 also formed a hydrogen bond to M87 W92 of the major helix. Lastly, C4BPα1 Ser42 was proximal to M87 D94 of the minor helix, and again with slight rearrangement could form a hydrogen bond to this amino acid.

The contribution of M87 amino acids to C4BP-binding was evaluated by Ala- or other-substitution mutagenesis. Intact M87 protein was used, and binding of His-tagged versions of wild-type or Ala-substituted M87 protein to intact C4BP was quantified by ELISA. Relative to wild-type M87 protein, Y81A, E89A, and D96A bound C4BP ∼21-fold less well on average, which indicated that the hydrophobic contact to C4BP R64 and polar contacts to K63 and R39, respectively, were crucial (Fig. 2D). In M2 protein, the hydrophobic contact to the C4BPα1 Arg39 nook is essential (via F75) (10), but here C4BP-binding decreased only 2-fold for M87 W92A. The M87 W92R substitution had a more dramatic effect, reducing C4BP-binding 22-fold, likely due to Arg-Arg charge repulsion. Contact to C4BPα2 R64 by M87 E85 was intermediate in importance, as the M87 E85A substitution resulted in only a 6-fold decrease in C4BP-binding.

The two contacts from the minor M87 helix, D80 and D94, were even less consequential, with C4BP-binding in D80A unchanged and in D94A decreased only 2-fold. Loss of C4BP-binding for intact M87 Y81A, E85A, E89A, and D96A was not due to loss of structure or stability, as evaluated by circular dichroism spectra and melting curves of these mutant proteins compared to wild-type intact M87 protein (Supp. Fig. S2). M87 W92R was previously shown to have the same structure and stability as wild-type M87 protein (22).

We next scored a collection of 179 M protein sequences for patterns that matched the C4BP-binding motif identified structurally and functionally for M87 protein. Sequences that had identical or chemically similar amino acids corresponding to the essential M87 amino acids Y81, E85, E89, and D96 were scored highly, while sequences lacking an identical or chemically similar amino acid at any of these positions were eliminated. Ala-substitution of M87 W92 and D94 had an intermediate effect on C4BP-binding, and thus sequences with identical or chemically similar amino acids at these positions were scored positively but not as highly as the essential M87 amino acids. Other amino acids were tolerated at these latter positions, except for positively charged ones at the position corresponding to W92, and hydrophobic ones that cannot form hydrogen bonds at the position corresponding to D94. While M87 D80 appeared not to influence C4BP-binding, this amino acid was located next to the important C4BPα2 amino acid R64, and thus sequences with a positively charged amino acid at this position were eliminated. The proper heptad position of these amino acids was also evaluated by scoring the *a* and *d* positions of three consecutive heptads. However, due to the often irregular heptads of M proteins (23, 24), only four *a* or *d* positions out of six in the three heptads needed to have coiled coil-stabilizing amino acids (25, 26). We noticed that this scoring scheme produced a rank-ordered list that included M24 protein near the bottom (Supp. Table S2). As the M24 strain does not bind C4BP (9), we used the score for M24 protein as a cut-off and included only M proteins that scored higher. This cut-off resulted in the identification of 19 M proteins, including M87, as having an M87 pattern for C4BP-binding (Fig. 2E and Supp. Table S2). M proteins are assigned to two clades, X and Y, with C4BP-binding assorting to clade X (27). Eleven of the 19 M proteins in the M87 pattern belong to a strain that binds C4BP, with all 11 belonging to clade X (9), whereas the other eight have not been evaluated for C4BP-binding, with most of these (6 of 8) belong to clade X.

### M68-C4BP interactions

The structure of the complex between M68^N100^ and C4BPα1-2 was determined to 2.5 Å resolution limit by SAD phasing using SeMet-labeled M68^N100^ (Supp. Table S1). The architecture of the complex differed substantially from that of M87 protein and those observed previously (10). M68^N100^ formed an α-helical dimer but not a coiled coil (Fig. 3A). Nearly the full extent of M68^N100^ was visualized in one of the helices (aa 42-135, ∼140 Å long), albeit with some breaks in main chain density, while the other helix spanned aa 52-127 without any breaks. The two M68 helices were asymmetrically disposed and mostly parallel to one other, except for the very N-terminal segment (aa 42-77), where one of the helices wrapped around the other. About two heptads of this N-terminal segment (aa 58-70) bound C4BP. Only a single C4BPα1-2 molecule was evident, and in this molecule the C4BPα1 domain was largely disordered. Both M68 helices contacted C4BPα2, with 1330 Å^2^ of surface area being buried, which is typical of M protein HVR-C4BP complexes (10). The shape complementarity between the two proteins was 0.58, which is also typical (10).

**Figure 3.**
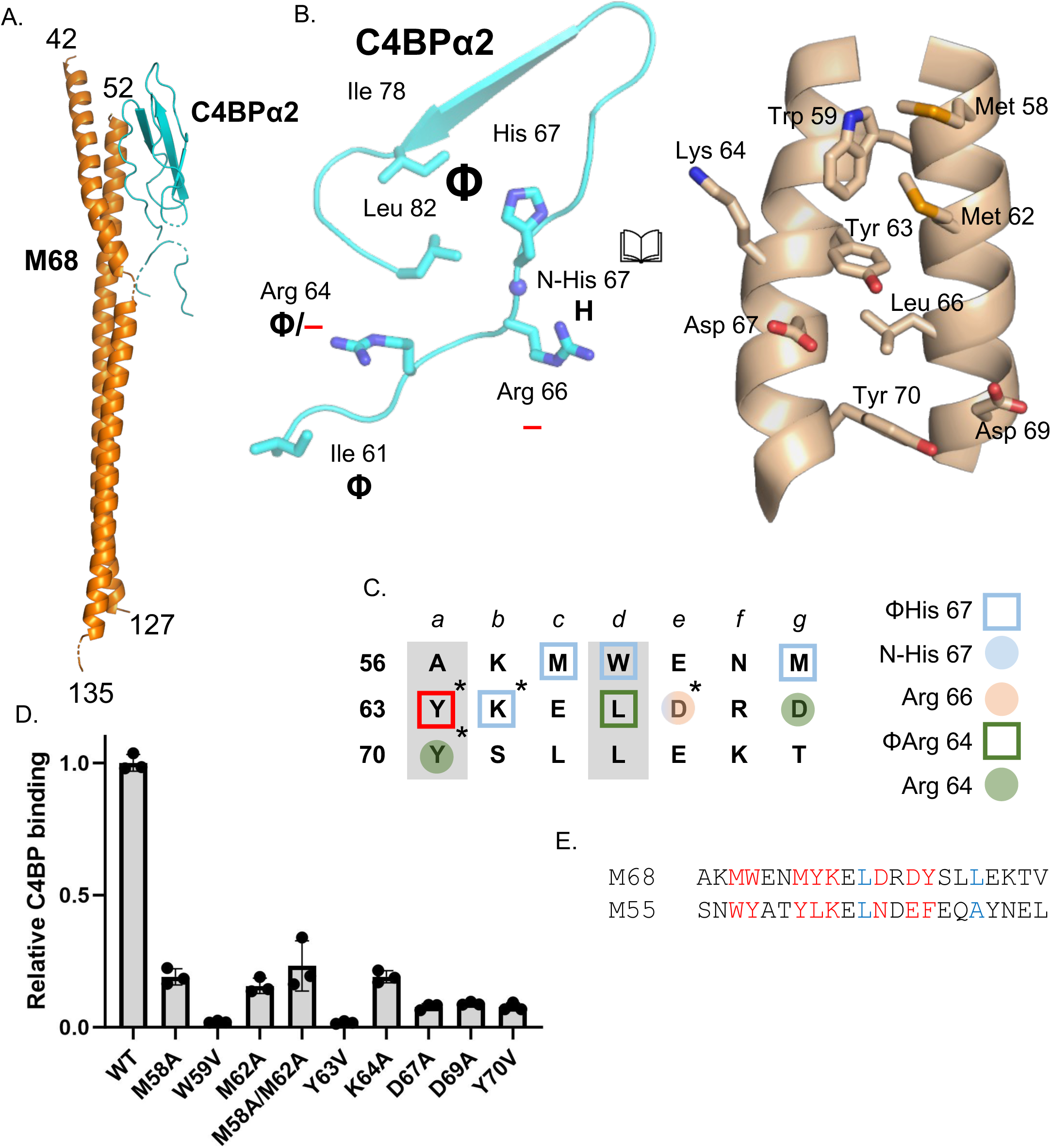
M68-C4BP interactions. **A.** Structure of M68 (orange) in complex with C4BPα2 (cyan), in cartoon representation. **B.** Interaction between C4BPα1-2 (left) and M68 (right), in cartoon (main chain) or bonds (side chains) representation, shown in open book format. Carbons are cyan for C4BPα1-2 and wheat for M87; for both, nitrogens are blue, and oxygens red. Chemical character of interactions denoted as follows: Φ, hydrophobic; —, negative; H, hydrogen bond. **C.** Heptad register of M68 56-76 (*a* and *d* positions in gray columns). Amino acids marked without an asterisk belong to one helix, and those with an asterisk to the second helix. Shapes indicate type of contact (square for hydrophobic, Φ; circle for polar), and coloring indicates which C4BP amino acid (depicted in legend to the right) was contacted. **D.** Binding of soluble, intact wild-type and mutant His_6_-M68 protein to immobilized intact C4BP, as evaluated by ELISA. Bound His_6_-M68 protein was detected with HRP-conjugated anti-His_6_ antibodies. Values were normalized by the value of C4BP-binding to wild-type His_6_-M68. Means and standard deviations from three biological replicates carried out in triplicate are shown. **E.** M protein sequences belonging to the M68 pattern of C4BP binding. Heptad positions are indicated above the sequences. Red indicates an observed or potential C4BP-binding amino acid, and blue a *d* heptad position.

The C4BPα2 quadrilateral contacted M68 protein, although with some variations (Figs. 3B and C). The C4BPα2 quadrilateral hydrophobic pocket (ΦHis 67), instead of contacting a single hydrophobic M protein amino acid (10), contacted a cluster: M68 Met 58, Trp 59, and Met 62 from one helix and Tyr 63, along with the alkyl chain of K64, from the second. The N-His 67 site formed a hydrogen bond with M68 D67 from the second helix. This same M87 amino acid formed a salt bridge with C4BPα2 R66. The other Arg in the quadrilateral, C4BPα2 R64, formed several interactions: a salt bridge to M68 D69 from the first helix, a hydrophobic contact to L66 from the first helix, and a π-cation interaction with M68 Y70 from the second helix. M68 Y70 also made a hydrophobic contact with C4BPα2 Ile61. Notably, all of the aromatic C4BP-contacting amino acids of M87 (W59, Y63, Y70) occupied the *a* or *d* heptad position (Fig. 3C).

The contribution of M68 amino acids to C4BP-binding was evaluated using intact M68 protein and intact C4BP, as described above for M87 protein. The contacts to the ΦHis 67 hydrophobic site were especially important. Valine-substitution of M68 W59, or Y63 reduced C4BP-binding ∼50-fold relative to wild-type M68 protein (Fig. 3D). Valine was chosen since these amino acids occupied the *a* or *d* heptad position. Valine is a stabilizing amino acid at the core *a* and *d* coiled-coil positions, whereas Ala is not (25, 26). Ala-substitution of Met 58, Met 62, or Lys 64, which also contacted the ΦHis 67 hydrophobic pocket reduced C4BP-binding by ∼6-fold. Likewise, dual-Ala substitution of M87 Met 58 and Met 62 reduced C4BP binding by ∼4-fold. Ala- or Val-substitution of M68 amino acids that contacted C4BPα2 R66 (M68 D67A) or R64 (M87 D69A and Y70V) reduced C4BP-binding by ∼12-fold. As validation of the structure of this unusual complex, we examined the structure and stability of M68 M58A/M62A and D67A, and found that their decreased C4BP-binding was a direct functional effect rather than an indirect effect on structure or stability (Supp. Fig. S3).

As for M87 protein, we scored M protein sequences for a pattern of C4BP-contacting amino resembling that of M68 protein, and for coiled-coil stabilizing amino acids at most *a* and *d* positions. Notably, M68 has three aromatic amino acids that are significant for C4BP-binding (W59, Y63, and Y70) at *a* or *d* positions. No other M protein in the set of 179 matched this pattern. If the restriction that these positions are occupied by only aromatic acids is relaxed to include hydrophobic ones larger than valine, then M55 protein is included in having an M68 pattern (Fig. 3E and Supp. Table S2). An M55 Strep A strain binds C4BP (9). M68 protein belongs to C4BP-binding clade X, whereas M55 protein is assigned to clade Y but is an outlier in this scheme (27). These results indicate that the unusual mode of interaction between C4BP and M68 protein is rare among M proteins.

### M2 and M22 Patterns

While contributions of M2 amino acids to C4BP-binding have been evaluated by mutagenesis previously (10), this prior study used a co-precipitation assay, unlike the ELISA used here for M87 and M68 proteins. To provide a common means for comparison, we examined C4BP-binding of Ala-substitution mutants of M2 protein by ELISA. We focused on three amino acids whose substitutions by Ala only slightly decreased C4BP-binding in the co-precipitation assay (10). These were M2 H61, which contacts the ΦHis 67 hydrophobic pocket; D62, which forms a hydrogen bond to N-His 67; and E68, which forms a salt bridge with C4BPα2 R64 (Fig. 4A). Ala-substitution of these amino acids in intact M2 protein led to ∼10-30-fold loss of C4BP-binding compared to intact wild-type M2 protein (Fig. 4B). A number of M2 amino acids were previously shown by the co-precipitation assay to be significant for C4BP-binding (10). These are F75, which contacts the C4BPα1 Arg39 nook, and E76 and D79, which form salt bridges with the C4BPα1 Arg39 nook, with only one or the other of these salt bridges being required (Fig. 4A). We constructed a scoring scheme for the M2 pattern based on these results, in which we also included the possibility of hydrophobic contacts to C4BP2α R64, as seen in M49 protein (10). This scoring scheme yielded a rank-ordered list that had M12 protein at the bottom (Supp. Table S2). As the M12 strain does not bind C4BP (9), we used the score for the M12 protein as a threshold. This resulted in the identification of 17 M proteins, including M2 and M49, that have the M2 motif (Fig. 3C and Supp. Table S2). Of these, 9 are from Strep A strains shown to bind C4BP, with all 9 belong to clade X (9), whereas the others have not been tested, with more than half of these (5 of 8) belonging to clade X (27).

**Figure 4.**
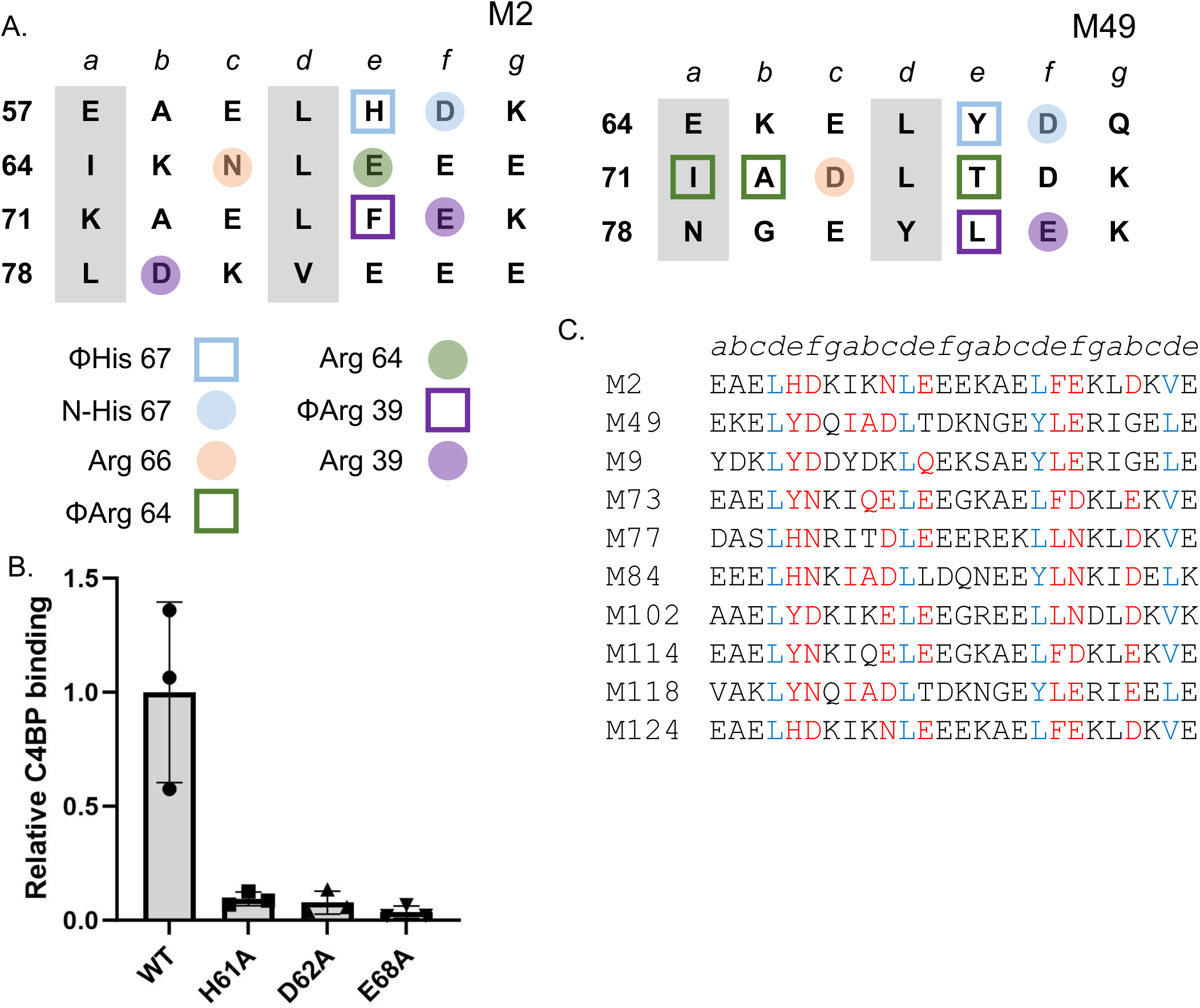
M2-C4BP interactions. **A.** Heptad register of (left) M2 (aa 57-84) and (right) M49 (aa 64-84) (*a* and *d* positions in gray columns). Shape indicates type of contact (square for hydrophobic, Φ; circle for polar), and coloring indicates which C4BP amino acid (depicted in legend at bottom) was contacted. **B.** Binding of soluble, intact His_6_-M2 protein constructs to immobilized intact C4BP evaluated by ELISA. Bound His_6_-M2 protein was detected with HRP-conjugated anti-His_6_ antibodies. Values were normalized by the value of C4BP-binding to wild-type His_6_-M2. Means and standard deviations from three biological replicates carried out in triplicate are shown. **C.** Partial list of M protein sequences belonging to the M2 pattern of C4BP binding. Heptad positions are indicated above the sequences. Red indicates an observed or potential C4BP-binding amino acid, and blue a *d* heptad position.

The M22 pattern has not been probed by mutagenesis (Fig. 5A), and therefore we examined binding by wild-type or mutant intact M22 protein to intact C4BP by ELISA. Ala-substitution of five M22 amino acids decreased C4BP-binding substantially (Fig. 5B). The largest decrease was ∼50-fold for M22 Y77A, which makes a hydrophobic contact to the C4BPα1 Arg39 nook (10). Ala-substitution of M22 I59, which makes hydrophobic contact to the C4BPα2 ΦHis 67 site, or M2 N60, which forms a hydrogen bond to C4BPα2 His 67, decreased C4BP-binding by ∼26-fold. Ala-substitution of M22 E65, which salt bridges with C4BPα2 R64, also decreased C4BP-binding substantially (10-fold), whereas Ala-substitution of M22 D64, which salt bridges with C4BPα2 R66, decreased it only slightly (∼3-fold). In M28 protein, the salt bridge to C4BPα2 R66 is made by a Glu in the following heptad (Fig. 5A, M22 E70). The equivalent position in M22 protein is also a Glu (E67). The M22 E67A substitution decreased C4BP-binding only slightly (∼3-fold), just like M22 D64A, but the dual substitution M22 D64A/E67A decreased C4BP-binding more substantially (∼13-fold), indicating that these two negatively charged M22 amino acids serve redundant roles. As above, we constructed a scoring scheme for the M22 pattern based on these results, and also took into consideration that the hydrophobic contact to the C4BPα1 Arg39 nook could occur from a *b* heptad position as in M28 protein, rather than an *e* heptad position as in M22 protein (Fig. 5A). As with the M2 pattern, M12 appeared near the bottom of the rank-ordered list, and the score for M12 protein was used as a threshold (Supp. Table S2). This resulted in 49 M proteins, including M22 and M28, with an M22 pattern (Fig. 5C and Supp. Table S2). Twenty-three of these belong to M strains shown to bind C4B, with all but one belonging to clade X (9), whereas the others have not been tested, with almost all (20 of 26) belonging to clade X (27).

**Figure 5.**
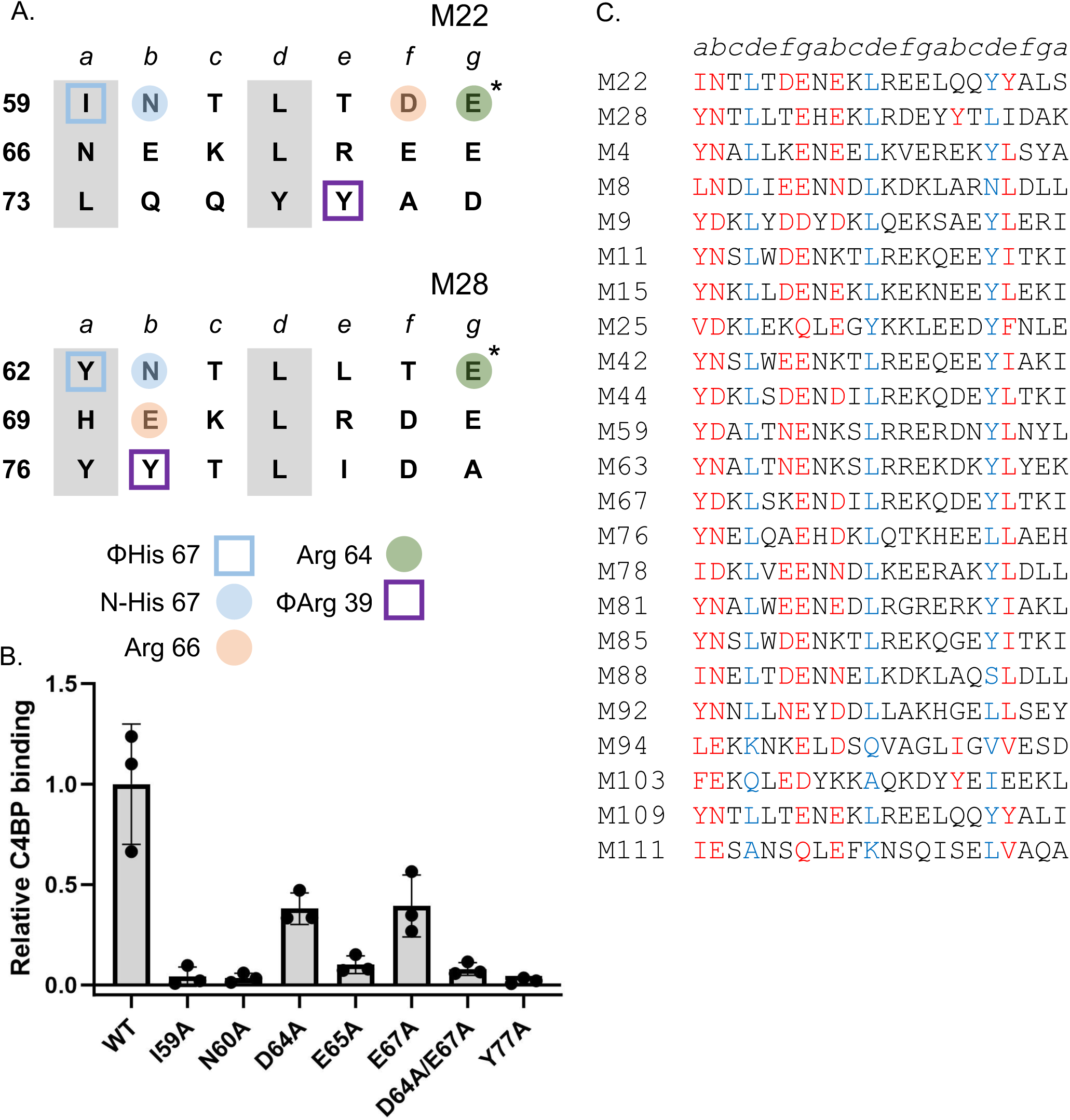
M22-C4BP interactions. **A.** Heptad register of (top) M22 (aa 59-79) and (bottom) M28 (aa 62-82) (*a* and *d* positions in gray columns). Amino acids marked with an asterisk belong to the minor helix, and those with an asterisk to the major helix. Coloring indicates which C4BP amino acid (depicted in legend at bottom) was contacted, and shape indicates type of contact (square for hydrophobic, Φ; circle for polar). **B.** Binding of soluble, intact His_6_-M22 protein constructs to immobilized intact C4BP evaluated by ELISA. Bound His_6_-M22 protein was detected with HRP-conjugated anti-His_6_ antibodies. Values were normalized by the value of C4BP-binding to wild-type His_6_-M22. Means and standard deviations from three biological replicates carried out in triplicate are shown. **C.** Partial list of M protein sequences belonging to the M22 pattern of C4BP binding. Heptad positions are indicated above the sequences. Red indicates an observed or potential C4BP-binding amino acid, and blue a *d* heptad position.

### C4BP binding by Enn proteins

The M2, M22, M68, or M87 patterns defined by experimental measurements still left about half of the 90 Strep A strains shown to bind C4BP unaccounted for (9). The genomes of most these strains encode an M-like Enn protein downstream of *emm*. Enn proteins are predicted to form α-helical coiled coils, and are sequence variable but to a lesser extent than M proteins (28, 29). Thus, we asked whether Enn proteins in such strains have C4BP-binding patterns. We identified an M2 pattern in 13 Enn proteins from 9 different M type strains, an M22 pattern in 70 Enn proteins from 45 strains, and an M87 pattern from 30 Enn proteins in 20 strains (Figs. 6A-C and Supp. Tables S3 and S4). The M2 pattern was found mainly in sub-group 5 (SG5) of Enn proteins (58%) and the M87 pattern mainly in SG1 (77%), while the M22 pattern was found across multiple sub-groups (Supp. Table S3) (30). The M68 pattern continued to be rare, with no Enn proteins identified to have this pattern. To validate these observations, we expressed and purified intact Enn300, which is encoded in an M18.6 strain and has the M22 pattern, and Enn292, which is encoded by an M91 strain and can be assigned to either M2 or M22 patterns (Fig. 6D). Both intact Enn300 and En292 bound intact C4BP (Fig. 6E), while the corresponding M proteins did not (M18.6) or bound weakly (M91). Ala-substitution of Enn300 N62 (equivalent to M22 N60), which is predicted to form a hydrogen bond to N-His 67, reduced C4BP binding by 4-fold relative to wild-type Enn300. Ala-substitution of Enn292 D70, which is predicted to likewise form a hydrogen bond to N-His 67 in the M2 pattern or instead contact C4BPα2 Arg 66 in the M22 pattern, reduced C4BP binding by 7-fold relative to wild-type Enn292. These results directly demonstrated C4BP-binding in Enn proteins, and validated the assignment of C4BP-binding patterns to Enn proteins.

**Figure 6.**
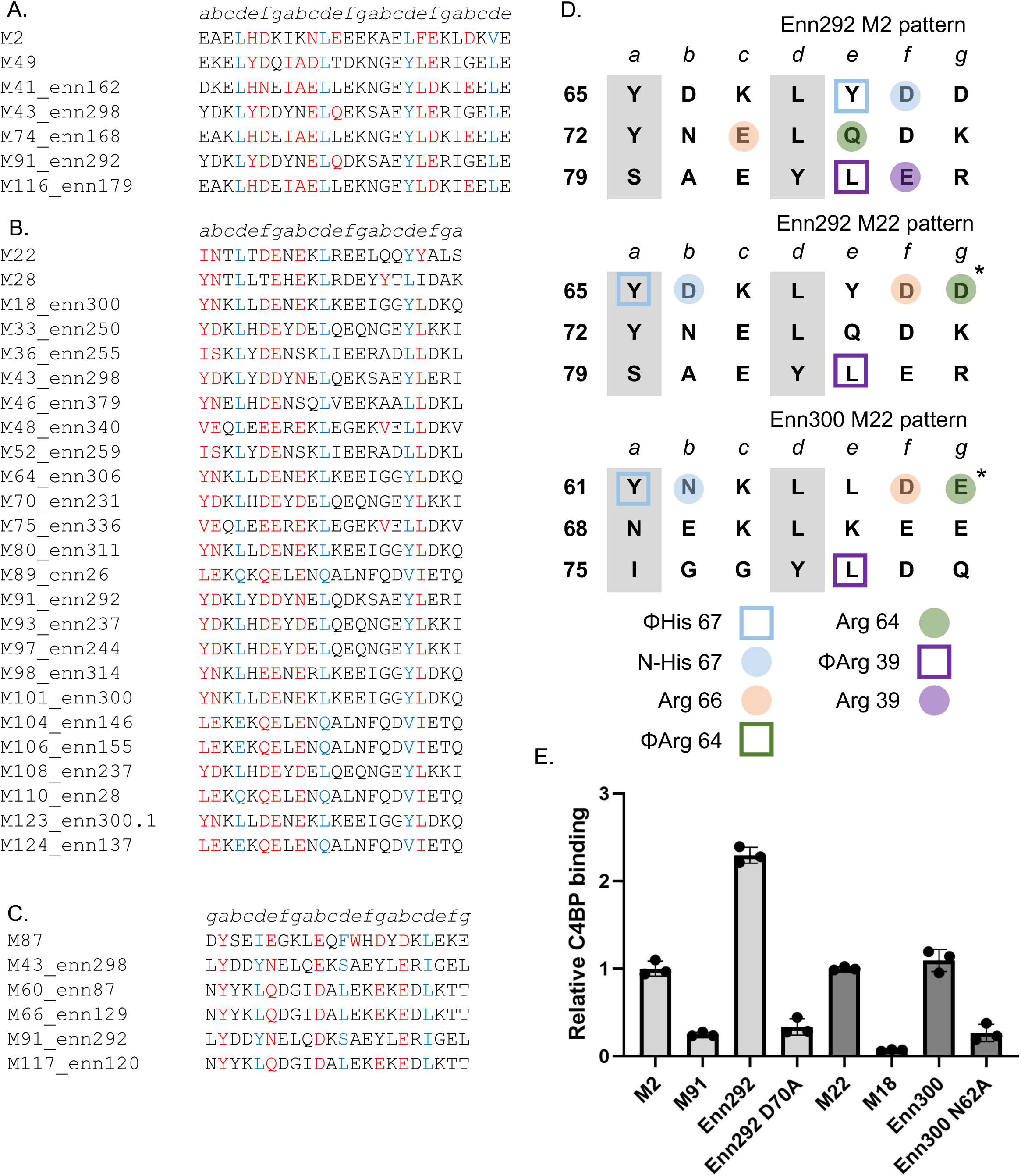
Enn protein-C4BP interactions. **A.** Enn protein sequences belonging to the M2 pattern (M2 and M49 sequences shown at top). Heptad positions are indicated above the sequences. Red indicates an observed or potential C4BP-binding amino acid, and blue a *d* heptad position. **B.** Same as panel A, except for the M22 pattern (M22 and M28 sequences shown at top). **C.** Same as panel A, except for the M87 pattern (M87 sequence shown at top). **D.** Heptad register of Enn292 (aa 64-84) in the M2 and M22 patterns, and Enn300 (aa 60-80) in the M22 pattern (*a* and *d* positions in gray columns). Amino acids marked with an asterisk belong to the major helix, and those without an asterisk to the minor helix. Shape indicates type of contact (square for hydrophobic, Φ; circle for polar), and coloring indicates which C4BP amino acid (depicted in legend to the right) is predicted to be contacted. **E.** Binding of soluble, intact His_6_-M or -Enn protein constructs (wild-type or Ala-substituted) to immobilized intact C4BP evaluated by ELISA. Bound His_6_-M or -Enn protein was detected with HRP-conjugated anti-His_6_ antibodies. Values were normalized by the value of C4BP-binding to wild-type His_6_-M2 protein for Enn292 and wild-type His_6_-M22 protein for Enn300. Means and standard deviations from three biological replicates carried out in triplicate are shown.

## Discussion

We sought to understand whether C4BP-binding patterns beyond the M2 and M22 patterns existed in M proteins. This was because somewhat less than half of 90 Strep A strains shown to bind C4BP have an M protein that could be assigned to either of these patterns (9, 10). To address this, several M proteins lacking an M2 or M22 pattern were identified to bind C4BP, and structures of the HVRs of two of these, M87 and M68, bound to a fragment of C4BP were determined. Neither M87 nor M68 HVRs precisely recapitulated the mode of C4BP interaction observed previously (10). For the M87 HVR, the C4BPα2 quadrilateral was not utilized, except for one amino acid from this grouping (R64), while the C4BPα1 Arg39 nook formed hydrophobic and polar contacts similar to ones observed previously (10). In contrast, the M68 HVR utilized the C4BPα2 quadrilateral, albeit using multiple amino acids rather than a single one to contact the ΦHis 67 site, but lacked contact to the C4BPα1 Arg39 nook, with the C4BPα1 domain in this complex almost entirely disordered.

Remarkably, in M87 protein the same site used for binding C4BP is also used for binding the human antimicrobial peptide LL-37 (22). Three M87 amino acids in particular form similar contacts to C4BP and LL-37. These are M87 Y81, E85, and W92, with the last of these forming hydrophobic contacts to an Arg in both C4BP (R39) and LL37 (R23). However, the structure of M87 protein differs considerably between these two complexes. With LL-37, the two M87 α-helices splay apart and no longer form a coiled coil in order to accommodate α-helical LL-37, which inserts between the two M87 helices, whereas with C4BP, the two M87 α-helices formed a coiled coil (Supp. Fig. S4). M87 in its unbound state also formed a coiled coil in this region (Supp. Fig. S4). Fourteen other M proteins have a sequence similar to the LL-37-binding motif of M87 (22), and all but one of these were identified to have a C4BP-binding pattern (Supp. Table S2). As expected, most have the M87 pattern, but the M22 and M68 patterns were also represented. These results point to the striking ability of a short coiled-coil sequence to have multiple functions through conformational dynamics.

The structures of M87 and M68 HVRs bound to C4BPα1-2 were used to guide mutational analysis to identify which M protein amino acids were most significant for binding. Based on these data, we searched M proteins for sequence patterns matching the C4BP-binding patterns of M87 and M68 proteins. Where appropriate, we used as a threshold the scores for M12 and M24 proteins, as these Strep A strains do not bind C4BP or have negligible binding (9). Numerous M proteins, 19 in all, have the M87 pattern, whereas the M68 pattern was rare, limited to 2 M proteins. We also carried out mutational analysis on M2 protein, taking into account its structure and that of the M49 HVR bound to C4BPα1-2, in addition to prior mutagenesis results on M2 protein (10). We also carried out mutational analysis on M22 protein, taking into account its structure and that of the M28 HVR bound to C4BPα1-2. Pattern searches based on these data revealed that the M22 pattern was the most populated, shared in 49 M proteins. The M2 pattern was also well populated, with 17 M proteins. Some M proteins had more than one sequence segment compatible with a single C4BP-binding pattern (e.g., two segments of M81 have an M22 pattern), and four M proteins (i.e. M8, M9, M81 and M103) had sequences compatible with two patterns (Supp. Table S2). Even with cutoffs applied, M3 protein from the very low C4BP-binding M3 strain was identified to have sequences compatible with M22 and M87 patterns (9). M3 protein has an unusually low coiled-coil propensity (31), even for an M protein (23, 24), and the absence of coiled-coil structure may impede M3 from binding C4BP. In all, 79 M protein types, excluding M3, out of 179 were identified as having an M2, M22, M68, or M87 C4BP-binding pattern.

M-like proteins also contribute to C4BP-binding. Deletion of the M-like Enn protein in M18 and M53 strains result in loss of C4BP-binding to the bacterial surface (32, 33). Complementation to restore C4BP-binding and verify the role of the Enn protein was not carried out in these studies. We demonstrated direct interaction between Enn300 protein, which is expressed by the M18 strain and has an M22 pattern, with C4BP. We also validated the M22 pattern in Enn300 through mutagenesis. In the M53 strain (specifically AP53), expression of the Enn protein requires inactivation of CovRS (33), a two-component system that negatively controls a multitude of virulence factors (34). The AP53 strain expresses Enn254, which we identified has an M22 pattern (Supp. Table S3). In the study that identified C4BP-binding in 90 strains, the M53 strain failed to bind C4BP (9), presumably due to lack of expression of the Enn protein. As with M proteins, some Enn proteins have at least two different types of C4BP-binding patterns (Fig. 6D, Enn292).

The C4BP-binding patterns were largely or exclusively near the N-terminal ends of M and Enn proteins (95% and 100%, respectively), a location that presumably enables access for multi-chained C4BP (α_7_β_1_, and with protein S also bound) (35). The exceptions were a few M proteins which had the M22 pattern (e.g., M221, aa 193). Further investigation is required to determine whether C4BP can access a C-terminal site closer to the bacterial cell wall. The most N-terminally located pattern was that of M77 protein (at amino acid 49, just 7 amino acids from the N-terminus), and in a previous study we found that M77 protein, despite having an excellent M2 patter, does not bind C4BP in a statistically significant manner *in vitro* (21). The very N-terminal portions of M proteins tend to lack coiled-coil character, which is true of M77 protein, and thus insufficient structuring may explain the lack of C4BP-binding. Structuring of the coiled coil may be enhanced by the binding of another protein, as has been shown for the interaction of the M-like protein H (PrtH) with C4BP requiring simultaneous binding of the Fc domain of IgG to PrtH (36). PrtH lacks a known C4BP-binding pattern.

In all, we identified C4BP-patterns in 111 Strep A strains of differing M types (Supp. Table S4, excluding M3). These comprise 71 of the 90 strains shown to bind C4BP (9). Among these 71 strains, 33 have both an M and an Enn protein with a C4BP-binding pattern, 9 only an M protein with a C4BP-binding pattern, and 29 only an Enn protein with a C4BP-binding pattern. Three strains that do not bind or bind C4BP only weakly (i.e., M24, M39, M53) have an Enn protein with a C4BP-binding pattern. As observed for the AP53 strain (33), interaction with C4BP in these strains may require mutation of CovRS or another regulatory system for expression of the Enn protein. In addition, 37 strains that to our knowledge have not been tested for C4BP-binding have a C4BP-binding pattern in their M proteins; Enn proteins were not investigated in these strains. In summary, our results indicate that C4BP-binding patterns are prevalent in Strep A strains of differing M types, being present in their M or Enn proteins, or frequently both, and provide further motivation for development of conserved C4BP-binding 3D patterns as vaccine immunogens.

## Experimental Methods

### DNA manipulation

Coding sequences for M and Enn proteins were cloned into a modified pET28b vector (Novagen) that contained sequences encoding an N-terminal His_6_-tag followed by a PreScission protease cleavage site. Amino acid substitutions and deletions were introduced with the QuickChange II Site-Directed mutagenesis kit (Stratagene), according to the manufacturer’s directions.

### Protein expression and purification

Expression of M proteins, Enn proteins, and C4BPα1-2 in *Escherichia coli* Gold (DE3) BL21 and purification of these proteins were carried out as previously described (10), with minor modifications. M and Enn protein constructs composed of their N-terminal 100 amino acids were purified only on Ni^2+^-NTA resin (Thermofisher Scientific) with the size exclusion chromatography step omitted, followed by dialysis against 20 mM Tris-HCl, pH 8.0, 100 mM NaCl. Constructs used for crystallization, namely C4BPα1-2, M87^N100^, and M68^N100^, were additionally subjected to PreScission protease (GenScript) digestion according to manufacturer’s instructions to remove the His_6_-tag. Intact M and Enn proteins were purified by affinity chromatography on Ni^2+^-NTA resin followed by size exclusion chromatography on a Superdex 200 16/60 column that had been pre-equilibrated with 20 mM Tris-HCl, pH 8.0, 100 mM NaCl. Fractions containing pure, non-degraded proteins of interest were pooled.

Selenomethionine (SeMet) labeling of M68^N100^ was performed using SelenoMet Medium (Molecular Dimensions) through feedback inhibition of methionine biosynthesis, as previously described (37). SeMet-M68^N100^ was purified as above, except buffers were supplemented with 5 mM DTT.

### Crystallization and data collection

SeMet-M68^N100^ and M87^N100^ were mixed with C4BPα1-2 in equimolar ratio. The buffer for the M68^N100^ mixture was exchanged by ultrafiltration through a 3,500 MWCO membrane (Millipore; 4,500 x *g*, 30 min, 4 °C) to 20 mM Tris-HCl, pH 8.0, 100 mM NaCl, and samples were concentrated to 10 mg/ml by ultrafiltration through a 3,500 MWCO membrane. The buffer for the M87^N100^ mixture was exchanged the same way to 10 mM Tris-HCl, pH 8.0, and samples were concentrated to 15 mg/ml, as described above. All crystallization trials were carried out at 20 °C using the hanging drop vapor diffusion method.

For crystallization of SeMet-M68^N100^-C4BPα1-2, 1 µl of protein sample was mixed with 1 µl of precipitant solution (100 mM MES-NaOH, pH 6.0, 18% PEG 3350, 10% ethylene glycol, 0.2 M CaCl_2_). Small (50 × 10 × 10 µm^3^) crystals, which grew in 2-7 days, were crushed and diluted 100-fold with precipitant solution to constitute a seed stock. The seed stock (0.2 µl) was mixed with 0.9 µl of SeMet-M68^N100^-C4BPα1-2 and 0.9 µl of 100 mM MES-NaOH, pH 6.0, 14% PEG 3350, 10% ethylene glycol, 0.2 M CaCl_2_, 1 mM DTT. Single crystals (300 x 50 x 50 μm^3^), which grew after 4-7 days, were transferred to cryoprotectant composed of 100 mM MES-NaOH, pH 6.0, 14% PEG 3350, 30% ethylene glycol, 0.2 M CaCl_2_, and 1 mM DTT, and flash cooled in liquid N_2_ prior to data collection. Diffraction data were collected at SSRL (beamline 9-2) at 0.9789, 0.9184, and 0.9793 Å wavelength (inflection, high energy, and remote, respectively), integrated with XDS (38), and scaled with Aimless (39).

For co-crystallization of M87^N100^-C4BPα1-2, 1 µl of protein sample was mixed with 1 µl of precipitant solution (0.1 M sodium citrate, pH 4.8, 0.2 M KH_2_PO_4_, 25% w/v PEG 3350). Crystals (400 x 80 x 80 μm^3^), which grew after about 7 days, were cryoprotected with precipitant solution supplemented with 20% glycerol, and flash frozen in liquid N_2_. Diffraction data were collected at APS (beamline 24-ID-E) at 0.979 Å, and automatically processed using Rapid Automated Processing of Data (RAPD), including XDS for integration and Aimless for scaling.

### Structure determination and refinement

Phase determination by multiwavelength anomalous dispersion, phase refinement, and initial model building for SeMet-M68^N100^-C4BPα1-2 were carried out with AutoBuild from the Phenix suite (40). The model was modified using Coot, as guided by the inspection of σA-weighted 2Fo-Fc and Fo-Fc omit maps. Multiple cycles of building followed by refinement were performed using Refine (41) from the Phenix suite with default settings. At late stages TLS parameters were introduced, with each chain constituting a single TLS group. Side chains with no electron density beyond Cβ were truncated at Cβ.

Phases for the M87^N100^-C4BPα1-2 complex were determined by molecular replacement using Phaser (42), with C4BPα1-2 (PDB 5I0Q) as the search model. The model was built manually into well-defined electron density using Coot, and modified as guided by the inspection of σA-weighted 2Fo-Fc and Fo-Fc omit maps. Refinement was performed using Refine from the Phenix suite with default settings and TLS parameters. Each chain constituted a single TLS group. Side chains with no electron density beyond Cβ were truncated at Cβ.

Figures of molecular models were generated with PyMol (The PyMOL Molecular Graphics System, Version 2.5.5 Schrödinger, LLC). Interaction interfaces between M proteins and C4BP were analyzed with PISA (43).

### CD spectroscopy

CD spectra were measured on a J-1500 Circular Dichroism Spectrometer (Jasco). Samples were exchanged by dialysis through a 10,000 MWCO membrane into 5 mM sodium phosphate, pH 7.9, and diluted to ∼0.125-0.250 mg/ml. Wavelength spectra were recorded between190-260 nm at 37 °C with a 0.5 nm interval. Thermal melting was carried out between 20-75 °C in 1 °C increments, with the CD signal being monitored at 222 nm. Wavelength scans and thermal melts were carried out in duplicate, and the data were averaged.

### ELISA

ELISAs were performed as previously described (21). In short, high-binding 96-well microtiter plates (Corning) were coated with intact C4BP (Athens Research and Technology; 5 µg/ml) in 100 μl PBS, pH 7.4 overnight at 4 °C. Wells were incubated with 100 μl of purified N-terminally His_6_-tagged M or Enn proteins (final concentration of 10 µg/ml). HRP-conjugated mouse anti-His antibody (BioLegend) diluted 1:5000 in 0.1% BSA/TBST was used for detection. Background from binding to uncoated wells was subtracted.

### Sequence scoring

Files containing 179 M protein or 143 Enn protein sequences (from NCBI GenBank) (29) were searched for C4BP-binding patterns. The sequences were in fasta format, and a python script was written to carry out the search. Sequences of 23-27 amino acids in length (i.e., more than three heptads but less than four) were searched. Only sequences that had coiled-coil stabilizing amino acids (i.e., V, L, I, M, F, or Y) (25, 26) at four or more *a* or *d* positions in three consecutive heptads were considered.

For the M87 pattern, at the position equivalent to M87 D80, the occurrence of a K or R eliminated the sequence from consideration (by giving this position a score of -100; only sequences that had a total score >0 were considered). At the position equivalent to M87 Y81, the occurrence of a I, L, M, F, Y, or W resulted in a score of +1. The occurrence of any other amino acid at this position resulted in elimination of the sequence. This was true for all positions at which specific amino acids were not excluded, except if stated otherwise. At positions equivalent to M87 E85, E89, and D96, the occurrence of a D, E, N, or Q resulted in a score of +1 for the first two and +0.7 for the last two; this scoring scheme was used at most positions occupied by D, E, N, or Q, and was called ‘DENQ’ scoring. At the position equivalent to M87 W92, the occurrence of a K or R eliminated the sequence, but the occurrence of a V, L, I, M, F, Y, or W (the term ‘hydrophobic amino acid’ refers to this set below) resulted in a score of +0.5. At the position equivalent to M87 D94, the occurrence of L, I, M, or F eliminated the sequence, but the occurrence of D or E resulted in a score of +0.5, N or Q a score of +0.3, and S or T a score of +0.1.

For the M68 pattern, at the positions equivalent to M68 M58 and M62, the occurrence of a hydrophobic amino acid resulted in a score of +1. At the positions equivalent to M68 W59, Y63, and Y70, the occurrence of I, L, M, F, Y, or W resulted in a score of +1. At the position equivalent to M68 K64, the occurrence of K or R resulted in a score of +1. At positions equivalent to M68 D67 and D69, DENQ scoring was used.

For the M2 pattern, the scoring was as follows. At the position equivalent to M2 H61, the occurrence of a His or a hydrophobic amino acid resulted in a score of +1. At the position equivalent to M2 D62, DENQ scoring was used, in addition to scoring Ser or Thr at this position with +0.5. At the position equivalent to M2 N66, we used the observation that if the preceding position were occupied by a Lys (equivalent to M2 K65), then the occurrence of an Asp or even Ala is preferred over Asn (10). Thus, if the position equivalent to M2 K65 were occupied by a Lys or Arg, then if the position equivalent to M2 N66 were occupied by a D or E, this resulted in a score of +1, whereas if it were occupied by an N or Q, this resulted in a score of +0.5. In the absence of a positively charged amino acid at the preceding position, DENQ scoring was applied. At the position equivalent to M2 E68, DENQ scoring was used. Since the polar contact of M2 E68 with C4BPα2 R64 is replaced in M49 with hydrophobic contacts to the alkyl chain of R64, the following was taken into account. In case the position equivalent to M2 E68 was not occupied by D, E, N, or Q, then if in the following heptad the *a* position (equivalent to M49 I71) were occupied by a coiled coil-stabilizing amino acid, the *b* position (equivalent to M49 A72) by Ala, and the *e* position (equivalent to M49 T75) by a small amino acid that can form a hydrophobic contact (T, V, or L), this resulted in a score of +1. At the position equivalent to M2 F75, the occurrence of a hydrophobic amino acid resulted in a score of +1. As M2 E76 and D79 form redundant salt bridges to C4BPα1 R39, the occurrence of D or E at either position resulted in a score of +1. If neither position were occupied by D or E, then the occurrence of an N or Q at either position resulted in a score of +0.7.

For the M22 pattern, the position equivalent to M22 I59, the occurrence of a coiled-coil stabilizing amino acid resulted in a score of +1. At the position equivalent to M22 N60, DENQ scoring was used, in addition to scoring Ser or Thr at this position with +0.5. In M22, D64 (heptad position *f*) forms a salt bridge to C4BPα2 R66, whereas in M28, E70 (*b* heptad position, equivalent to M22 position 67) does. Thus, if a D, E, N, or Q occupied either the position equivalent of M22 positions 64 or 67, DENQ scoring was used. At the position equivalent to M22 E65, DENQ scoring was used. M22 Y77 (*e* heptad position) forms a hydrophobic contact to C4BPα1 R39, whereas M28 Y77 (*b* heptad position, equivalent to M22 position 74) does the same. Therefore, the occurrence of a hydrophobic amino acid at the equivalent of M22 position 74 or 77 resulted in a score of +1.

## Supporting Information

Supplementary Figures S1-S4.

Supplementary Tables S1-S4.

## Acknowledgements

We thank Naieemah Mershon for help with genetic construct preparation and protein purification. This work was supported by NIH 1R01AI154149 (PG) and the Belgian Fonds National de la Recherche Scientifique FNRS PDR N°0227.20 (AB and PS). Use of the Stanford Synchrotron Radiation Lightsource, SLAC National Accelerator Laboratory, is supported by the U.S. Department of Energy, Office of Science, Office of Basic Energy Sciences under Contract No. DE-AC02-76SF00515. The SSRL Structural Molecular Biology Program is supported by the DOE Office of Biological and Environmental Research, and by the National Institutes of Health, National Institute of General Medical Sciences (P30GM133894). The contents of this publication are solely the responsibility of the authors and do not necessarily represent the official views of NIGMS or NIH. This research used resources of the Advanced Photon Source, a U.S. Department of Energy (DOE) Office of Science user facility operated for the DOE Office of Science by Argonne National Laboratory under Contract No. DE-AC02-06CH11357.

## References

1. Barnett TC, Bowen AC, Carapetis JR. The fall and rise of Group A Streptococcus diseases. Epidemiol Infect. 2018:1–6. PubMed PMID: 30109840.

2. Nelson GE, Pondo T, Toews KA, Farley MM, Lindegren ML, Lynfield R, Aragon D, Zansky SM, Watt JP, Cieslak PR, Angeles K, Harrison LH, Petit S, Beall B, Van Beneden CA. Epidemiology of Invasive Group A Streptococcal Infections in the United States, 2005-2012. Clin Infect Dis. 2016;63(4):478–86. PubMed PMID: 27105747.

3. Carapetis JR, Beaton A, Cunningham MW, Guilherme L, Karthikeyan G, Mayosi BM, Sable C, Steer A, Wilson N, Wyber R, Zuhlke L. Acute rheumatic fever and rheumatic heart disease. Nat Rev Dis Primers. 2016;2:15084. PubMed PMID: 27188830.

4. Nitsche-Schmitz DP, Chhatwal GS. Host-pathogen interactions in streptococcal immune sequelae. Curr Top Microbiol Immunol. 2013;368:155–71. PubMed PMID: 23212184.

5. Carapetis JR, Steer AC, Mulholland EK, Weber M. The global burden of group A streptococcal diseases. Lancet Infect Dis. 2005;5(11):685–94. PubMed PMID: 16253886.

6. Berggard K, Johnsson E, Morfeldt E, Persson J, Stalhammar-Carlemalm M, Lindahl G. Binding of human C4BP to the hypervariable region of M protein: a molecular mechanism of phagocytosis resistance in Streptococcus pyogenes. Mol Microbiol. 2001;42(2):539–51. PubMed PMID: 11703674.

7. Carlsson F, Berggard K, Stalhammar-Carlemalm M, Lindahl G. Evasion of phagocytosis through cooperation between two ligand-binding regions in *Streptococcus pyogenes* M protein. J Exp Med. 2003;198(7):1057–68. PubMed PMID: 14517274.

8. Ermert D, Shaughnessy J, Joeris T, Kaplan J, Pang CJ, Kurt-Jones EA, Rice PA, Ram S, Blom AM. Virulence of group A *Streptococci* is enhanced by human complement inhibitors. PLoS Pathog. 2015;11(7):e1005043. PubMed PMID: 26200783.

9. Persson J, Beall B, Linse S, Lindahl G. Extreme sequence divergence but conserved ligand-binding specificity in *Streptococcus pyogenes* M protein. PLoS Pathog. 2006;2(5):e47. PubMed PMID: 16733543.

10. Buffalo CZ, Bahn-Suh AJ, Hirakis SP, Biswas T, Amaro RE, Nizet V, Ghosh P. Conserved patterns hidden within group A Streptococcus M protein hypervariability recognize human C4b-binding protein. Nat Microbiol. 2016;1:16155. PubMed PMID: 27595425.

11. McMillan DJ, Dreze PA, Vu T, Bessen DE, Guglielmini J, Steer AC, Carapetis JR, Van Melderen L, Sriprakash KS, Smeesters PR. Updated model of group A *Streptococcus* M proteins based on a comprehensive worldwide study. Clin Microbiol Infect. 2013;19(5):E222–9. PubMed PMID: 23464795.

12. Ghosh P. Variation, Indispensability, and Masking in the M protein. Trends Microbiol. 2018;26(2):132–44. PubMed PMID: 28867148.

13. Johnsson E, Thern A, Dahlback B, Heden LO, Wikstrom M, Lindahl G. A highly variable region in members of the streptococcal M protein family binds the human complement regulator C4BP. J Immunol. 1996;157(7):3021–9. PubMed PMID: 8816411.

14. Morfeldt E, Berggard K, Persson J, Drakenberg T, Johnsson E, Lindahl E, Linse S, Lindahl G. Isolated hypervariable regions derived from streptococcal M proteins specifically bind human C4b-binding protein: implications for antigenic variation. J Immunol. 2001;167(7):3870–7. PubMed PMID: 11564804.

15. Vekemans J, Gouvea-Reis F, Kim JH, Excler JL, Smeesters PR, O’Brien KL, Van Beneden CA, Steer AC, Carapetis JR, Kaslow DC. The Path to Group A Streptococcus Vaccines: World Health Organization Research and Development Technology Roadmap and Preferred Product Characteristics. Clin Infect Dis. 2019;69(5):877–83. PubMed PMID: 30624673.

16. Sandin C, Carlsson F, Lindahl G. Binding of human plasma proteins to *Streptococcus pyogenes* M protein determines the location of opsonic and non-opsonic epitopes. Mol Microbiol. 2006;59(1):20–30. PubMed PMID: 16359315.

17. Lannergard J, Gustafsson MC, Waldemarsson J, Norrby-Teglund A, Stalhammar-Carlemalm M, Lindahl G. The hypervariable region of *Streptococcus pyogenes* M protein escapes antibody attack by antigenic variation and weak immunogenicity. Cell Host Microbe. 2011;10(2):147–57. PubMed PMID: 21843871.

18. Jones KF, Fischetti VA. The importance of the location of antibody binding on the M6 protein for opsonization and phagocytosis of group A M6 streptococci. J Exp Med. 1988;167(3):1114–23. PubMed PMID: 2450950.

19. Pandey M, Ozberk V, Calcutt A, Langshaw E, Powell J, Rivera-Hernandez T, Ho MF, Philips Z, Batzloff MR, Good MF. Streptococcal Immunity Is Constrained by Lack of Immunological Memory following a Single Episode of Pyoderma. PLoS Pathog. 2016;12(12):e1006122. PubMed PMID: 28027314.

20. Dale JB, Beachey EH. Localization of protective epitopes of the amino terminus of type 5 streptococcal M protein. J Exp Med. 1986;163(5):1191–202. PubMed PMID: 2422314.

21. Wang KC, Kuliyev E, Nizet V, Ghosh P. A conserved 3D pattern in a Streptococcus pyogenes M protein immunogen elicits M-type crossreactivity. J Biol Chem. 2023;299(8):104980. PubMed PMID: 37390991.

22. Kolesinski P, Wang KC, Hirose Y, Nizet V, Ghosh P. An M protein coiled coil unfurls and exposes its hydrophobic core to capture LL-37. Elife. 2022;11. PubMed PMID: 35726694.

23. Nilson BH, Frick IM, Akesson P, Forsen S, Bjorck L, Akerstrom B, Wikstrom M. Structure and stability of protein H and the M1 protein from Streptococcus pyogenes. Implications for other surface proteins of gram-positive bacteria. Biochemistry. 1995;34(41):13688–98. PubMed PMID: 7577960.

24. Stewart CM, Buffalo CZ, Valderrama JA, Henningham A, Cole JN, Nizet V, Ghosh P. Coiled-coil destabilizing residues in the group A Streptococcus M1 protein are required for functional interaction. Proc Natl Acad Sci U S A. 2016;113(34):9515–20. PubMed PMID: 27512043.

25. Tripet B, Wagschal K, Lavigne P, Mant CT, Hodges RS. Effects of side-chain characteristics on stability and oligomerization state of a de novo-designed model coiled-coil: 20 amino acid substitutions in position “d”. J Mol Biol. 2000;300(2):377–402. PubMed PMID: 10873472.

26. Wagschal K, Tripet B, Lavigne P, Mant C, Hodges RS. The role of position a in determining the stability and oligomerization state of alpha-helical coiled coils: 20 amino acid stability coefficients in the hydrophobic core of proteins. Protein Sci. 1999;8(11):2312–29. PubMed PMID: 10595534.

27. Sanderson-Smith M, De Oliveira DM, Guglielmini J, McMillan DJ, Vu T, Holien JK, Henningham A, Steer AC, Bessen DE, Dale JB, Curtis N, Beall BW, Walker MJ, Parker MW, Carapetis JR, Van Melderen L, Sriprakash KS, Smeesters PR, Group MPS. A systematic and functional classification of *Streptococcus pyogenes* that serves as a new tool for molecular typing and vaccine development. J Infect Dis. 2014;210(8):1325–38. PubMed PMID: 24799598.

28. Whatmore AM, Kapur V, Musser JM, Kehoe MA. Molecular population genetic analysis of the enn subdivision of group A streptococcal emm-like genes: horizontal gene transfer and restricted variation among enn genes. Mol Microbiol. 1995;15(6):1039–48. PubMed PMID: 7623660.

29. Frost HR, Davies MR, Delforge V, Lakhloufi D, Sanderson-Smith M, Srinivasan V, Steer AC, Walker MJ, Beall B, Botteaux A, Smeesters PR. Analysis of Global Collection of Group A Streptococcus Genomes Reveals that the Majority Encode a Trio of M and M-Like Proteins. mSphere. 2020;5(1). PubMed PMID: 31915226.

30. Frost HR, Guglielmini J, Duchene S, Lacey JA, Sanderson-Smith M, Steer AC, Walker MJ, Botteaux A, Davies MR, Smeesters PR. Promiscuous evolution of Group A Streptococcal M and M-like proteins. Microbiology (Reading). 2023;169(1). PubMed PMID: 36748538.

31. Ludwiczak J, Winski A, Szczepaniak K, Alva V, Dunin-Horkawicz S. DeepCoil-a fast and accurate prediction of coiled-coil domains in protein sequences. Bioinformatics. 2019;35(16):2790–5. PubMed PMID: 30601942.

32. Perez-Caballero D, Garcia-Laorden I, Cortes G, Wessels MR, de Cordoba SR, Alberti S. Interaction between complement regulators and Streptococcus pyogenes: binding of C4b-binding protein and factor H/factor H-like protein 1 to M18 strains involves two different cell surface molecules. J Immunol. 2004;173(11):6899–904. PubMed PMID: 15557185.

33. Agrahari G, Liang Z, Mayfield JA, Balsara RD, Ploplis VA, Castellino FJ. Complement-mediated opsonization of invasive group A Streptococcus pyogenes strain AP53 is regulated by the bacterial two-component cluster of virulence responder/sensor (CovRS) system. J Biol Chem. 2013;288(38):27494–504. PubMed PMID: 23928307.

34. Graham MR, Smoot LM, Migliaccio CA, Virtaneva K, Sturdevant DE, Porcella SF, Federle MJ, Adams GJ, Scott JR, Musser JM. Virulence control in group A Streptococcus by a two-component gene regulatory system: global expression profiling and in vivo infection modeling. Proc Natl Acad Sci U S A. 2002;99(21):13855–60. PubMed PMID: 12370433.

35. Dahlback B. Purification of human C4b-binding protein and formation of its complex with vitamin K-dependent protein S. Biochem J. 1983;209(3):847–56. PubMed PMID: 6223625.

36. Ermert D, Weckel A, Magda M, Morgelin M, Shaughnessy J, Rice PA, Bjorck L, Ram S, Blom AM. Human IgG Increases Virulence of Streptococcus pyogenes through Complement Evasion. J Immunol. 2018;200(10):3495–505. PubMed PMID: 29626087.

37. Doublie S. Production of selenomethionyl proteins in prokaryotic and eukaryotic expression systems. Methods Mol Biol. 2007;363:91–108. PubMed PMID: 17272838.

38. Kabsch W. XDS. Acta Crystallogr D Biol Crystallogr. 2010;66(Pt 2):125–32. PubMed PMID: 20124692.

39. Evans PR, Murshudov GN. How good are my data and what is the resolution? Acta Crystallographica Section D. 2013;69(7):1204–14.

40. Liebschner D, Afonine PV, Baker ML, Bunkoczi G, Chen VB, Croll TI, Hintze B, Hung LW, Jain S, McCoy AJ, Moriarty NW, Oeffner RD, Poon BK, Prisant MG, Read RJ, Richardson JS, Richardson DC, Sammito MD, Sobolev OV, Stockwell DH, Terwilliger TC, Urzhumtsev AG, Videau LL, Williams CJ, Adams PD. Macromolecular structure determination using X-rays, neutrons and electrons: recent developments in Phenix. Acta Crystallogr D Struct Biol. 2019;75(Pt 10):861–77. PubMed PMID: 31588918.

41. Afonine PV, Grosse-Kunstleve RW, Echols N, Headd JJ, Moriarty NW, Mustyakimov M, Terwilliger TC, Urzhumtsev A, Zwart PH, Adams PD. Towards automated crystallographic structure refinement with phenix.refine. Acta Crystallographica Section D. 2012;68(4):352–67.

42. McCoy AJ, Grosse-Kunstleve RW, Adams PD, Winn MD, Storoni LC, Read RJ. Phaser crystallographic software. Journal of Applied Crystallography. 2007;40:658–74.

43. Krissinel E, Henrick K. Inference of macromolecular assemblies from crystalline state. J Mol Biol. 2007;372(3):774–97. PubMed PMID: 17681537.

