## Supplementary Information for "Conservation of C4BP-binding Sequence Patterns in *Streptococcus pyogenes* M and Enn Proteins"

**Supplementary Figures S1-S4**

**Supplementary Tables S1-S4**

### Supplementary Figure Legends

**Figure S1. Structure of M87$^{N100}$ and its complex with C4BPα1-2. A.** Superposition of the two M87 α-helices (orange and wheat, except 80-96 in gray). **B.** Superposition of interfaces between M87$^{N00}$ (aa 79-96) and C4BPα1-2, based on the position of C4BPα1-2. One interface has M87$^{N100}$ in orange and C4BPα1-2 in cyan, and the other has M87$^{N100}$ in wheat and C4BPα1-2 in pale cyan.

**Figure S2. Structure and stability of wild-type and mutant M87 proteins.**

**A.** Circular dichroism (CD) spectra of intact wild-type and mutant M87 proteins at 25 °C. The data are an average of two independent measurements. **B.** Thermal melting curves of intact wild-type and mutant M87 proteins, as monitored by the CD mean molar residue ellipticity at 222 nm. The data are an average of two independent measurements.

**Figure S3. Structure and stability of wild-type and mutant M68 proteins.**

**A.** CD spectra of intact wild-type and mutant M68 proteins at 25 °C. The data are an average of two independent measurements. **B.** Thermal melting curves of intact wild-type and mutant M68 proteins, as monitored by the CD mean molar residue ellipticity at 222 nm. The data are an average of two independent measurements.

**Figure S4. Comparison of C4BP-binding region in different structural states of M87.** The ligand-binding site of M87 is shown in cartoon representation, with M87 in its complex with C4BPα1-2 in magenta and with LL-37 in cyan. M87 in its uncomplexed form is in green.

Figure S1

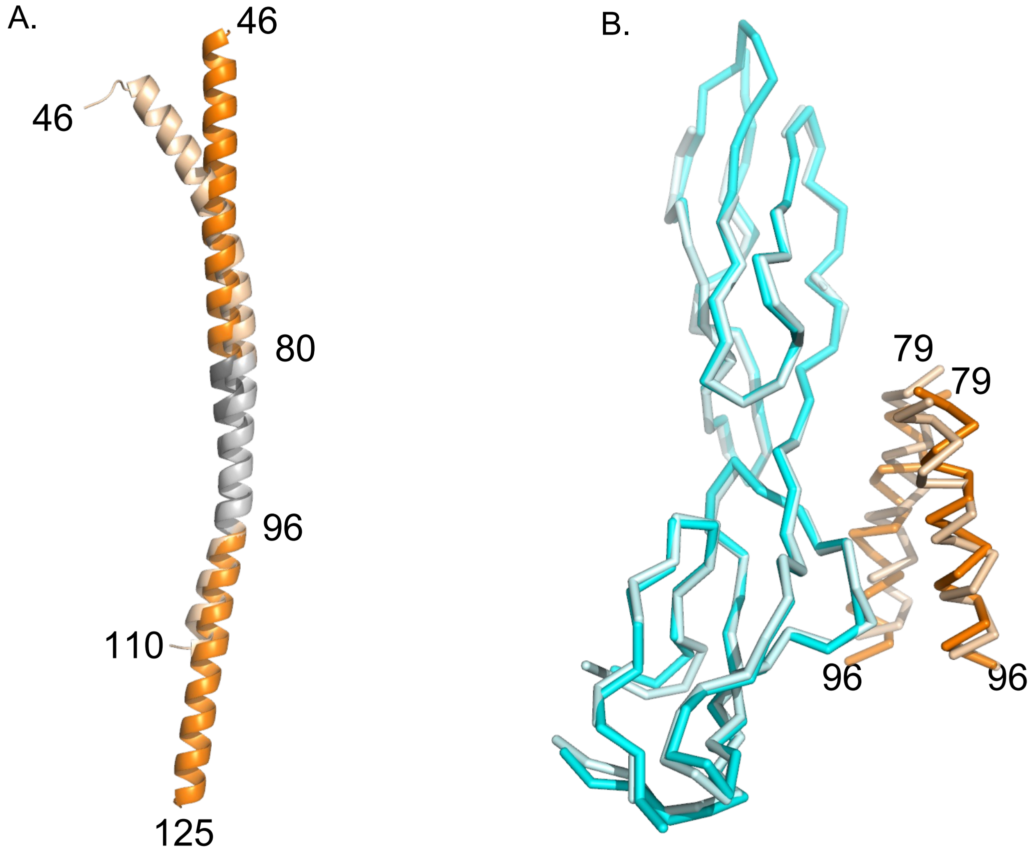

Figure S2

A.

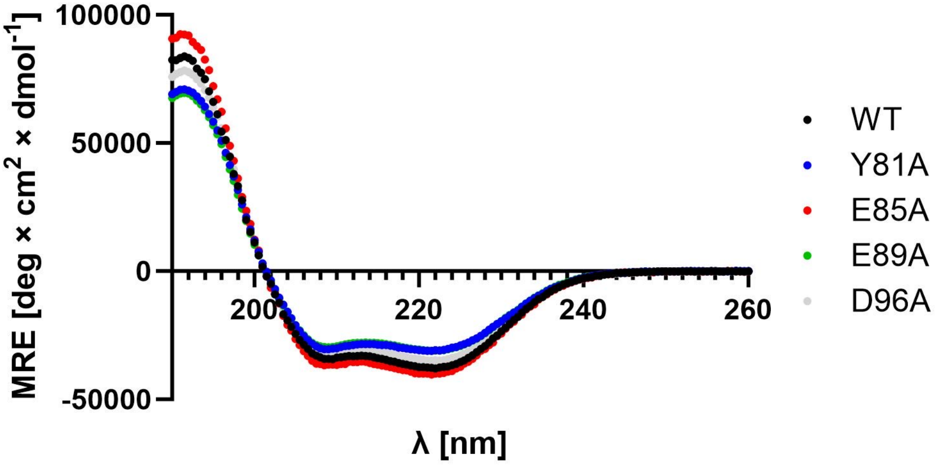
100000
50000
0
-50000
MRE [deg × cm2 × dmol-1]
200
220
240
260
λ [nm]
WT
Y81A
E85A
E89A
D96A

B.

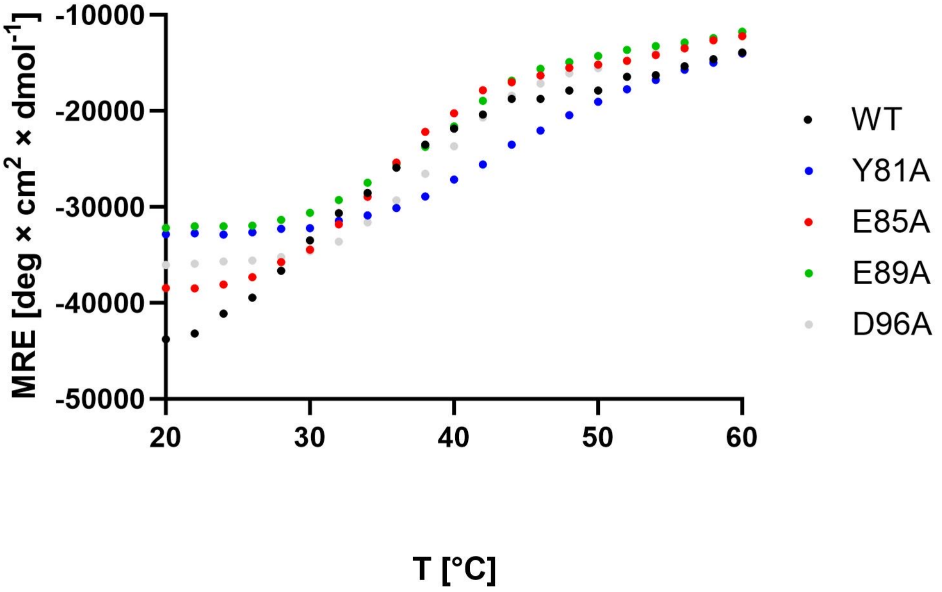
-10000
-20000
-30000
-40000
-50000
MRE [deg × cm2 × dmol-1]
20
30
40
50
60
T [°C]
WT
Y81A
E85A
E89A
D96A

Figure S3

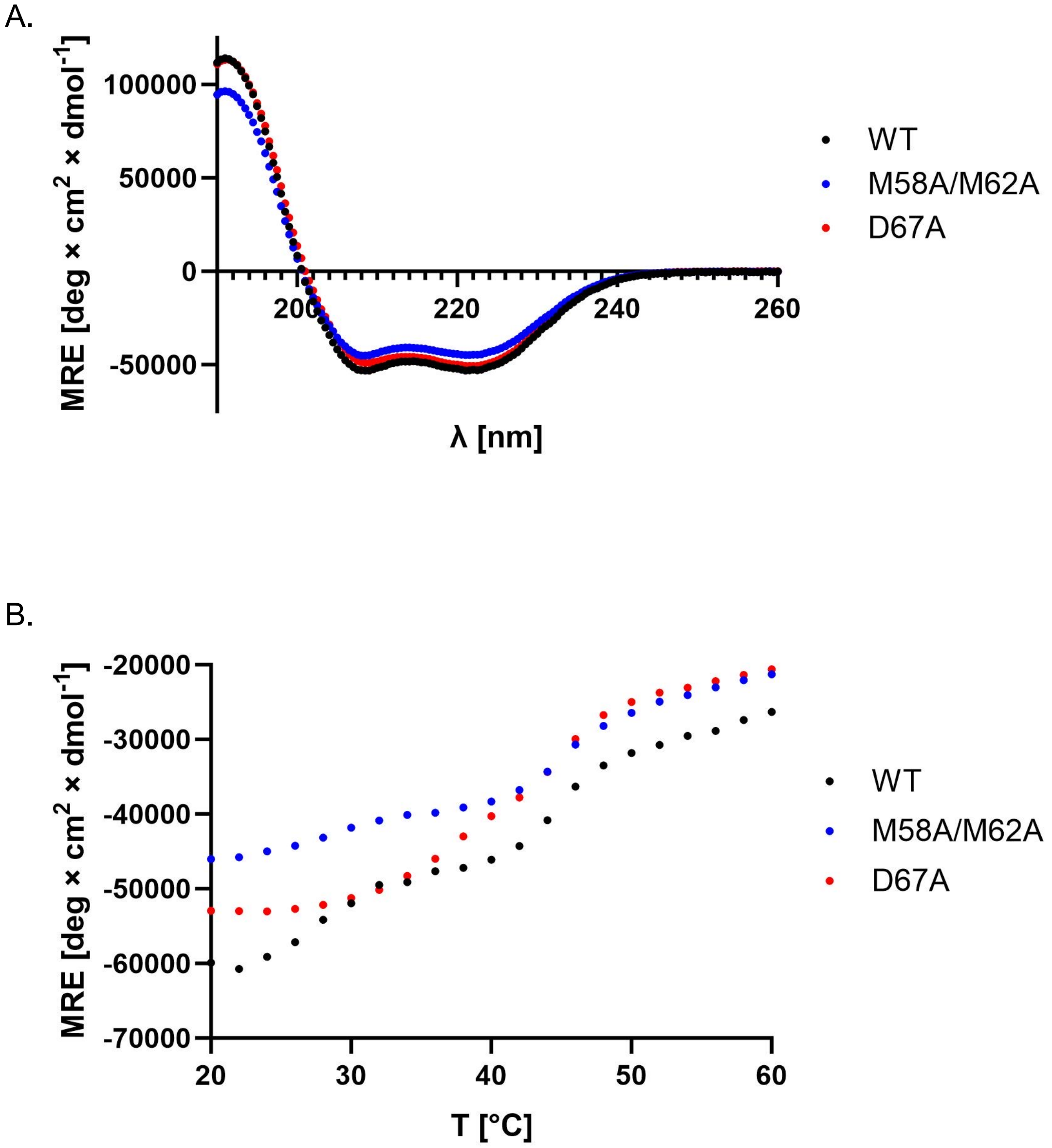
A.
MRE [deg × cm2 × dmol-1]
100000
50000
0
-50000
200
220
240
260
λ [nm]
WT
M58A/M62A
D67A
B.
MRE [deg × cm2 × dmol-1]
-20000
-30000
-40000
-50000
-60000
-70000
20
30
40
50
60
T [°C]
WT
M58A/M62A
D67A

Figure S4

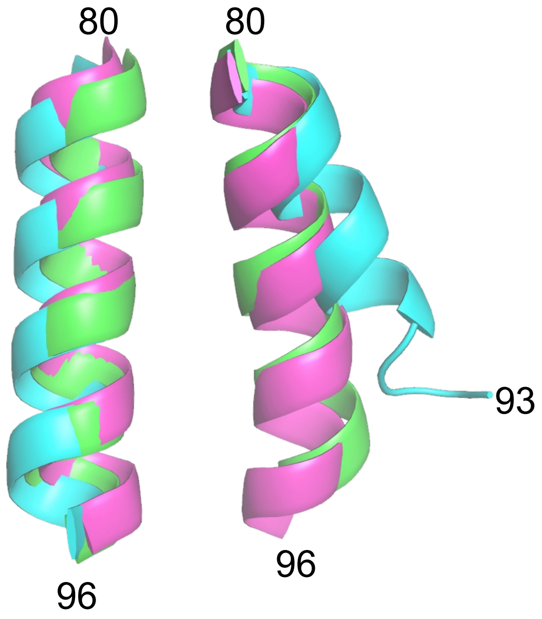

**Table S1. Crystallographic Data Collection and Model Refinement**

| | M87$^{N100}$/C4BPα1-2 | SeMet-M68$^{N100}$/C4BPα1-2 |
|---|---|---|
| **Data Collection** | | |
| Wavelength (Å) | 0.98 | 0.98 |
| Resolution range (Å) | 68.81 - 2.69 (2.82 – 2.69) | 39.30 – 2.50 (2.60 – 2.50) |
| Space group | P 2 $2_1$ $2_1$ | P $2_1$ $2_1$ 2 |
| Cell dimensions | | |
| a, b, c (Å) | 43.18 73.80 190.40 | 107.04 49.66 80.40 |
| α, β, γ (°) | 90.0 90.0 90.0 | 90.0 90.0 90.0 |
| Unique reflections | 17478 (2045) | 15422 (1715) |
| Multiplicity | 6.8 (3.0) | 13.1 (13.2) |
| Completeness (%) | 98.64 (87.13) | 99.80 (99.80) |
| I/σ(I) | 20.3 (1.4) | 8.7 (0.8) |
| Wilson B-factor (Å$^2$) | 54.92 | 43.96 |
| $R_{merge}$ | 0.09 (0.953) | 0.275 (3.524) |
| $CC_{1/2}$ | 0.997 (0.452) | 0.997 (0.650) |
| **Refinement** | | |
| Resolution range (Å) | 68.81 - 2.69 (2.82 – 2.69) | 37.63 – 2.50 (2.60 – 2.50) |
| No. of reflections (work/test set) | 17414/872 | 13083/1327 |
| $R_{work}$/$R_{free}$ | 0.24/0.28 (0.35/0.37) | 0.31/0.34 (0.39/0.46) |
| No. of non-hydrogen atoms | 2705 | 1632 |
| macromolecules | 2675 | 1606 |
| ligands | 1 | 6 |
| solvent | 29 | 20 |
| R.m.s. deviations | | |
| bonds (Å) | 0.004 | 0.002 |
| angles (°) | 0.62 | 0.34 |
| Ramachandran plot | | |
| favoured (%) | 97.50 | 97.71 |
| outliers (%) | 0.00 | 0.00 |
| Rotamer outliers (%) | 0.84 | 0.00 |
| Clashscore | 4.11 | 2.51 |
| Average B-factor (Å$^2$) | 56.45 | 62.58 |
| macromolecules | 56.59 | 62.70 |
| ligands | 72.13 | 52.32 |
| solvent | 43.52 | 64.67 |
| Number of TLS groups | 4 | 3 |
| PDB code | 8TCB | 8TGT |

**Supplementary Table S2. C4BP-binding Patterns in M proteins.**
List of sequences of M proteins sequences with C4BP-binding patterns. Sequences are ordered by their score, followed by the M type number. Red amino acids are observed or predicted to bind C4BP. Start indicates sequence number (including the signal sequence). M proteins that belong to strains identified as binding C4BP (1) are indicated with bold font. * indicates an M protein shown or predicted to bind LL-37 (2). A horizontal line, if present, indicates a threshold set by an M protein from a strain that does not bind C4BP (e.g., M24 or M12) (1).

### M87 Pattern

| M | Start | Sequence | Score | Clade |
|---|---|---|---|---|
| **87*** | **80** | **DYSEIEGKLEQFWHDYDKLEKE** | **5** | **X** |
| **112** | **59** | **EIAALQEEKEKLLNELDNVKEE** | **4.7** | **X** |
| 31 | 98 | YLKVLNDWAEKLLKELNGDDVK | 4.4 | Y |
| **79*** | **79** | **DYSQIEEKLEQFGHDYDKLEKE** | **4.5** | **X** |
| **103** | **83** | **DYYEIEEKLEQFGRDYDKLEKE** | **4.5** | **X** |
| **107*** | **86** | **GYFNLEDRIEKLGSDYEKLEKE** | **4.5** | **X** |
| 144* | 97 | GYFNLEDRIEKLGSDYEKLEKE | 4.5 | X |
| 231 | 91 | GYFNLEDRIEKLGSDYEKLEKE | 4.5 | X |
| 3 | 96 | YLKGLNDWAERLLQELNGEDVK | 4.4 | X |
| **81** | **83** | **ELNEKNRKLEADIADLQDVIED** | **4.4** | **X** |
| 133 | 62 | YLKGLNDWAERLLQELNGEDVK | 4.4 | Y |
| **8** | **62** | **LIEENNDLKDKLARNLDLLDNT** | **4.3** | **X** |
| **9** | **62** | **LYDDYDKLQEKSAEYLERIGEL** | **4.3** | **X** |
| **82*** | **88** | **EYENLEETLQKFGNDYERLENK** | **4.2** | **X** |
| **90*** | **73** | **DYTVLQEAIEGISSENEKLKSE** | **4.2** | **X** |
| 168* | 75 | YYSLLQETIENMSLENENLDKL | 4.2 | X |
| 209* | 79 | DYSQIQEELEQFGHDYDKLEKE | 4.2 | X |
| **58*** | **80** | **NYETLEKGYDDLEKTLENFGES** | **4.1** | **X** |
| **58*** | **87** | **GYDDLEKTLENFGESYDKLENK** | **4.1** | **X** |
| 180* | 59 | DYEILEKRYDALEKTLENFGES | 4.1 | X |
| 31 | 63 | LLERVEQLDNSHNNNYEQYRHY | 4 | X |
| 34 | 63 | LYDDYNELQDKSAEYLERIGEL | 4 | X |
| 24 | 50 | TLEKVQERADKFEIENNTLKLK | 3.9 | Y |
| 88 | 52 | SISNNERLINELTDENNELKDK | 3.9 | X |
| 19 | 127 | QIQDLESQTQDLESQTQALESQ | 3.7 | Y |
| 76 | 58 | LYNELQAEHDKLQTKHEELLAE | 3.7 | X |
| 106 | 59 | IYDELQTKYDELQTKHEELLGE | 3.7 | X |
| 151 | 80 | ELEERQKNLEKLEHQFQVAADK | 3.7 | ND |

### M68 Pattern

| M | Start | Sequence | Score | Clade |
|---|---|---|---|---|
| **68*** | **56** | **AKMWENMYKELDRDYSLLEKTV** | **8** | **X** |
| **55** | **92** | **SNWYATYLKELNDEFEQAYNEL** | **7.7** | **Y** |

### M2 Pattern

| M | Start | Sequence | Score | Clade |
|---|---|---|---|---|
| **49** | **64** | **EKELYDQIADLTDKNGEYLERIGELE** | **6** | **X** |
| **102** | **52** | **AAELYDKIKELEEGREELLNDLDKVK** | **6** | **X** |
| 175 | 59 | EAELHDKIADLEEEKAELLNKLDKVE | 6 | X |
| 31 | 60 | IENLLERVEQLDNSHNNNYEQYRHYS | 5.7 | Y |
| 34 | 60 | YDKLYDDYNELQDKSAEYLERIGELE | 5.7 | X |
| **73** | **58** | **EAELYNKIQELEEGKAELFDKLEKVE** | **5.7** | **X** |

| | | | | |
|---|---|---|---|---|
| 77 | 49 | DASLHNRITDLEEEREKLLNKLDKVE | 5.7 | X |
| 84 | 56 | EEELHNKIADLLDQNEEYLNKIDELK | 5.7 | X |
| 114 | 65 | EAELYNKIQELEEGKAELFDKLEKVE | 5.7 | X |
| 118 | 56 | VAKLYNQIADLTDKNGEYLERIEELE | 5.7 | X |
| 137 | 58 | LYELYDEITNLEEERGKHLDRIEELE | 5.7 | X |
| 169 | 70 | HLDLIDNIREKDPQYRALMGENQDLR | 5.7 | X |
| 232 | 59 | EEKLHNKIADLLDQNEEYLNKIDELK | 5.7 | Y |
| 2 | 57 | EAELHDKIKNLEEEKAELFEKLDKVE | 5.5 | X |
| 124 | 65 | EAELHDKIKNLEEEKAELFEKLDKVE | 5.5 | X |
| 95 | 103 | WEQYINSYRDNNDRVQYYLEKANGTD | 5.4 | Y |
| 27 | 58 | YEELHTKYEELQTKHEELLGENDALK | 5.2 | X |

**M22 Pattern**

| M | Start | Sequence | Score | Clade |
|---|---|---|---|---|
| 9 | 59 | YDKLYDDYDKLQEKSAEYLERI | 5 | X |
| 11 | 93 | LDKKNKELDSRVTDLIDVIEHD | 5 | X |
| 34 | 60 | YDKLYDDYNELQDKSAEYLERI | 5 | X |
| 42 | 99 | LDKKNKELDSRVTDLIDVIGHD | 5 | X |
| 44 | 57 | YDKLSDENDILREKQDEYLTKI | 5 | X |
| 59 | 84 | LEKKNKELDSQVAGLIGVVESD | 5 | X |
| 63 | 82 | LEKKNKELDSQVAGLIGVVESD | 5 | X |
| 67 | 57 | YDKLSKENDILREKQDEYLTKI | 5 | X |
| 67 | 88 | LDKKNKELDSQVAGLIGVVESD | 5 | X |
| 78 | 55 | IDKLVEENNDLKEERAKYLDLL | 5 | X |
| 85 | 90 | LDKKNKELDSRVTDLIDVIEHD | 5 | X |
| 85 | 108 | IEHDDQELERKERMYEAFLKQS | 5 | X |
| 94 | 80 | LEKKNKELDSQVAGLIGVVESD | 5 | X |
| 103* | 70 | FEKQLEDYKKAQKDYYEIEEKL | 5 | X |
| 139 | 67 | YEKLWTEHEKLLGEHGELLGEH | 5 | X |
| 170 | 56 | VEQLEEEREKLEGEKVELLDKV | 5 | X |
| 172 | 65 | YESLYDEHEKLYDEKEKLYDEY | 5 | X |
| 174 | 53 | YDKLHDEYDELQEQNGEYLKKI | 5 | X |
| 182 | 106 | IEHDDQELERKGRMYEAFLKQS | 5 | X |
| 191 | 92 | LEKENKELDSQVTDLLDVIEHD | 5 | X |
| 236 | 56 | YDALRDENTGLRGDQKKLVKKL | 5 | X |
| 236 | 87 | LEKEKQELENQALNFQDVIETQ | 5 | X |
| 3 | 60 | IENLLDQVTQLYTKHNSNYQQY | 4.7 | X |
| 3 | 148 | YQDLDKDFDLAKQGYVLSDKRH | 4.7 | X |
| 4 | 59 | YNALLKENEELKVEREKYLSYA | 4.7 | X |
| 8 | 59 | LNDLIEENNDLKDKLARNLDLL | 4.7 | X |
| 11 | 62 | YNSLWDENKTLREKQEEYITKI | 4.7 | X |
| 11 | 111 | IEHDDQELKRKDRMYEAFLKQS | 4.7 | Y |
| 13 | 60 | YNKLSGDYDKLWDKHGELLSEY | 4.7 | X |
| 15 | 63 | YNKLLDENEKLKEKNEEYLEKI | 4.7 | X |
| 22 | 59 | INTLTDENEKLREELQQYYALS | 4.7 | X |
| 25* | 68 | VDKLEKQLEGYKKLEEDYFNLE | 4.7 | X |
| 28 | 62 | YNTLLTEHEKLRDEYYTLIDAK | 4.7 | X |
| 42 | 61 | YNSLWEENKTLREEQEEYIAKI | 4.7 | Y |
| 51 | 59 | LDELYNELHDENSQLVEEKAAL | 4.7 | X |
| 51 | 63 | YNELHDENSQLVEEKAALLDKL | 4.7 | X |
| 57 | 164 | YQNTKREYDLIEEELGKKLKEN | 4.7 | Y |
| 59 | 57 | YDALTNENKSLRERDNYLNYL | 4.7 | X |
| 76 | 59 | YNELQAEHDKLQTKHEELLAEH | 4.7 | X |
| 81 | 53 | YNALWEENEDLRGRERKYIAKL | 4.7 | X |
| 81 | 102 | IEDNDQEIKRKDRMYEAFLKQS | 4.7 | X |

| | | | | |
|---|---|---|---|---|
| **85** | **59** | **YNSLWDENKTLREKQGEYITKI** | **4.7** | **X** |
| **88** | **60** | **INELTDENNELKDKLAQSLDLL** | **4.7** | **X** |
| **92** | **57** | **YNNLLNEYDDLLAKHGELLSEY** | **4.7** | **X** |
| **109** | **54** | **YNTLLTENEKLREELQQYYALI** | **4.7** | **X** |
| **111** | **183** | **IESANSQLEFKNSQISELVAQA** | **4.7** | **Y** |
| 113 | 58 | YNKLSDERNNLLDQNGDLLDQN | 4.7 | X |
| 133 | 114 | YQDLDKDFDLAKQGYVLSDKRH | 4.7 | Y |
| 158 | 59 | YNKLSNERDNLLGENGKLWDEN | 4.7 | X |
| 165 | 60 | YNKLVEENSKLQKQLEEYLDSS | 4.7 | X |
| 176 | 66 | YNKLVDENDKLQKQLEEYLDSS | 4.7 | X |
| 177 | 57 | YDALTNENKSLRKERDNYLNYL | 4.7 | X |
| 193 | 57 | YDRLDEQNHKLVNDNHKLVNDK | 4.7 | Y |
| 205 | 61 | YNKLLDENEKLKEEIGGYLDKQ | 4.7 | X |
| 221 | 193 | IESANSQLEFKNSQISELVAQA | 4.7 | Y |
| 50 | 56 | ISKLYDENSKLIEERADLLGKL | 4.5 | X |
| 65 | 58 | YSKLLNENDILRDKQDDYLTKI | 4.5 | X |
| 95 | 154 | LSTLQEELKTAKSVYELAVLST | 4.5 | Y |
| 134 | 56 | ISKLYDENSKLIEERADLLDKL | 4.5 | X |
| 157 | 78 | ITDLTTELDEKVASFEAELGRN | 4.5 | Y |
| 183 | 55 | YSQLHDDYDKLQEQNGEYLKKI | 4.5 | X |
| 12 | 195 | VDETDRNLQQEKQKVLSLEQQL | 4.4 | Y |
| 49 | 50 | VEAAENNVSSVARREKELYDQI | 4.4 | X |
| 55 | 308 | LETINNNLLGNAKDMIAKLSAE | 4.4 | Y |
| 63 | 58 | YNALTNENKSLRREKDKYLYEK | 4.4 | X |
| 117 | 58 | YNELSGEYNKLLDQNGNLLDEN | 4.4 | X |
| 151 | 43 | VEVAENNVSSVARREKELYDQI | 4.4 | ND |
| 222 | 63 | LNNDLRQLEGKVRNLRSMMHEL | 4.4 | Y |
| 222 | 300 | LETINNNLLGNAKDMIAKLSAK | 4.4 | Y |
| 228 | 181 | VDETDRNLQQEKQKVLSLEQQL | 4.4 | Y |
| 229 | 197 | VDETDRNLQQEKQKVLSLEQQL | 4.4 | Y |
| 56 | 55 | LSELPGQAQAFSRAFLHEREKN | 3.9 | Y |
| 56.2 | 55 | LSELPGQAQAFSRAFLHERQKN | 3.9 | Y |

1. Persson J, Beall B, Linse S, Lindahl G. Extreme sequence divergence but conserved ligand-binding specificity in *Streptococcus pyogenes* M protein. PLoS Pathog. 2006;2(5):e47. PubMed PMID: 16733543.
2. Kolesinski P, Wang KC, Hirose Y, Nizet V, Ghosh P. An M protein coiled coil unfurls and exposes its hydrophobic core to capture LL-37. Elife. 2022;11. PubMed PMID: 35726694.

**Supplementary Table S3. C4BP-binding Patterns in Enn Proteins.**
List of sequences of Enn proteins sequences with C4BP-binding patterns. Sequences are indicated by the M strain followed by Enn protein identifier, and are ordered by the score followed by numerical order of the M type. Red amino acids are predicted to bind C4BP. Start indicates sequence number (including the signal sequence). Enn proteins that belong to strains identified as binding C4BP are indicated with bold font. A horizontal line, if present, indicates a cut-off threshold set by an M protein from a strain that does not bind C4BP (i.e., M24). Enn protein sub-groups (SG) are indicated.

#### M2 Pattern

| Enn | Start | Sequence | Score | SG |
|---|---|---|---|---|
| **M43_enn172** | **52** | **EAKLHDEIAELLEKNGEYLDKIEELE** | **6** | **SG5** |
| **M72_enn377** | **52** | **EAKLHDEIAELLEKNGEYLDKIEELE** | **6** | **SG6** |
| **M74_enn168** | **52** | **EAKLHDEIAELLEKNGEYLDKIEELE** | **6** | **SG5** |
| **M74_enn173** | **52** | **EAKLHDEIAELLEKNGEYLDKIEELE** | **6** | **SG5** |
| **M74_enn170** | **52** | **EAKLHDEIAELLEKNGEYLDKIEELE** | **6** | **SG5** |
| **M116_enn179** | **52** | **EAKLHDEIAELLEKNGEYLDKIEELE** | **6** | **SG5** |
| **M118_enn58** | **52** | **EAKLHDEIAELLEKNGEYLDKIEELE** | **6** | **SG3** |
| M24_enn282 | 58 | LYELYDEITNLEEERGKHLDRIEELE | 5.7 | SG5 |
| M39_enn283 | 58 | LYELYDEITNLEEERGKHLDRIEELE | 5.7 | SG5 |
| **M41_enn162** | **55** | **DKELHNEIAELLEKNGEYLDKIEELE** | **5.7** | **SG9** |
| **M41_enn167** | **56** | **DEELHNEIAELLEKNGEYLDKIEELE** | **5.7** | **SG8** |
| **M43_enn298** | **66** | **YDKLYDDYNELQEKSAEYLERIGELE** | **5.7** | **outlier** |
| **M91_enn292** | **65** | **YDKLYDDYNELQDKSAEYLERIGELE** | **5.7** | **outlier** |

#### M22 Pattern

| Enn | Start | Sequence | Score | SG |
|---|---|---|---|---|
| **M4_enn2** | **62** | **LEKQKQELENQALNFHDVIETQ** | **5** | **SG3** |
| **M4_enn14** | **56** | **YDALKDENTGLRGDQTKLVKKL** | **5** | **SG3** |
| **M4_enn16** | **56** | **YDTLKDENTGLRGDQTKLVKKL** | **5** | **SG3** |
| **M9_enn33** | **87** | **LEKQKQELENQALNFQDVIETQ** | **5** | **SG3** |
| **M15_enn36** | **87** | **LEKQKQELENQALNFQDVIETQ** | **5** | **SG3** |
| **M15_enn37** | **87** | **LEKQKQELENQALNFQDVIETQ** | **5** | **SG3** |
| **M22_enn151** | **93** | **LEKEKQELENQALNFQDVIETQ** | **5** | **outlier** |
| **M22_enn342** | **56** | **VEQLEEEREKLEGEKVELLDKV** | **5** | **SG8** |
| **M28_enn67** | **62** | **LEKQKQELENQALNFQDVIETQ** | **5** | **SG4** |
| **M33_enn250** | **53** | **YDKLHDEYDELQEQNGEYLKKI** | **5** | **SG5** |
| **M33_enn249** | **53** | **YDKLHDEYDELQEQNGEYLKKI** | **5** | **SG5** |
| **M33_enn248** | **53** | **YDKLHDEYDELQEQNGEYLKKI** | **5** | **SG5** |
| **M43_enn298** | **66** | **YDKLYDDYNELQEKSAEYLERI** | **5** | **SG9** |
| **M44_enn5** | **87** | **LEKQKQELENQALNFQDVIETQ** | **5** | **SG3** |
| **M44_enn11** | **87** | **LEKQKQELENQALNFQDVIETQ** | **5** | **SG3** |
| **M44_enn12** | **87** | **LEKQKQELENQALNFQDVIETQ** | **5** | **SG4** |
| **M48_enn340** | **56** | **VEQLEEEREKLEGEKVELLDKV** | **5** | **SG8** |
| **M49_enn40.1** | **56** | **YDALKDENTGLRGDQTKLVKKL** | **5** | **SG3** |
| **M49_enn40** | **56** | **YDALKDENTGLRGDQTKLVKKL** | **5** | **SG3** |
| M53_enn230 | 53 | YDKLHDEYDELQEQNGEYLKKI | 5 | SG5 |
| M53_enn234 | 53 | YDKLHDEYDELQEQNGEYLKKI | 5 | SG5 |
| **M58_enn48** | **56** | **YDALRDENTGLRGDRTKLVKKL** | **5** | **SG3** |
| **M58_enn51** | **56** | **YDALRDENTGLRGDRTKLVKKL** | **5** | **SG3** |
| **M58_enn52** | **56** | **YDALRDENTGLRGDRTKLVKKL** | **5** | **SG3** |
| **M59_enn240** | **53** | **YDKLHDEYDELQEQNGEYLKKI** | **5** | **SG5** |
| **M63_enn247** | **52** | **YDKLHDEYDELQEQNGEYLKKI** | **5** | **SG5** |
| **M68_enn10** | **87** | **LEKQKQELENQALNFQDVIETQ** | **5** | **SG3** |
| **M68_enn5.4** | **87** | **LEKQKQELENQALNFQDVIETQ** | **5** | **SG3** |

| | | | | |
|---|---|---|---|---|
| M70_enn231 | 53 | YDKLHDEYDELQEQNGEYLKKI | 5 | SG5 |
| M70_enn232 | 53 | YDKLHDEYDELQEQNGEYLKKI | 5 | SG5 |
| M73_enn5.1 | 87 | LEKQKQELENQALNFQDVIETQ | 5 | SG8 |
| M75_enn336 | 56 | VEQLEEEREKLEGEKVELLDKV | 5 | SG8 |
| M76_enn40 | 56 | YDALKDENTGLRGDQTKLVKKL | 5 | SG3 |
| M76_enn38 | 56 | YDALKDENTGLRGDQTKLVKKL | 5 | SG3 |
| M78_enn13 | 87 | LEKQKQELENQALNFQDVIETQ | 5 | SG3 |
| M78_enn68 | 87 | LEKQKQELENQALNFQDVIETQ | 5 | SG3 |
| M78_enn14 | 56 | YDALKDENTGLRGDQTKLVKKL | 5 | SG3 |
| M89_enn26 | 87 | LEKQKQELENQALNFQDVIETQ | 5 | SG3 |
| M90_enn3 | 87 | LEKQKQELENQALNFQDVIETQ | 5 | SG3 |
| M90_enn141 | 93 | LEKEKQELENQALNFQDVIETQ | 5 | SG2 |
| M91_enn292 | 65 | YDKLYDDYNELQDKSAEYLERI | 5 | SG8 |
| M93_enn237 | 53 | YDKLHDEYDELQEQNGEYLKKI | 5 | SG5 |
| M97_enn244 | 52 | YDKLHDEYDELQEQNGEYLKKI | 5 | SG5 |
| M102_enn152 | 90 | LEKEKQELENQALNFQDVIETQ | 5 | SG2 |
| M104_enn146 | 93 | LEKEKQELENQALNFQDVIETQ | 5 | SG2 |
| M106_enn155 | 90 | LEKEKQELENQALNFQDVIETQ | 5 | SG2 |
| M108_enn237 | 53 | YDKLHDEYDELQEQNGEYLKKI | 5 | SG5 |
| M109_enn17 | 56 | YDALKDENTGLRGDQTKLVKKL | 5 | SG4 |
| M110_enn28 | 87 | LEKQKQELENQALNFQDVIETQ | 5 | SG3 |
| M112_enn132 | 91 | LEKEKQELENQALNFQDVIETQ | 5 | SG2 |
| M114_enn22 | 87 | LEKQKQELENQALNFQDVIETQ | 5 | SG3 |
| M118_enn60 | 56 | YDALRDENTGLRGDQKKLVKKL | 5 | SG3 |
| M118_enn60 | 87 | LEKEKQELENQALNFQDVIETQ | 5 | SG3 |
| M118_enn23 | 87 | LEKQKQELENQALNFQDVIETQ | 5 | SG3 |
| M124_enn137 | 93 | LEKEKQELENQALNFQDVIETQ | 5 | SG2 |
| M124_enn138 | 93 | LEKEKQELENQALNFQDVIETQ | 5 | SG2 |
| M2_enn127 | 91 | LEAINKELNENYYKLQDGIDAL | 4.7 | SG1 |
| M18_enn300 | 61 | YNKLLDENEKLKEEIGGYLDKQ | 4.7 | SG9 |
| M46_enn379 | 59 | LDELYNELHDENSQLVEEKAAL | 4.7 | outlier |
| M46_enn379 | 63 | YNELHDENSQLVEEKAALLDKL | 4.7 | outlier |
| M64_enn306 | 61 | YNKLLDENEKLKEEIGGYLDKQ | 4.7 | SG9 |
| M80_enn311 | 61 | YNKLLDENEKLKEEIGGYLDKQ | 4.7 | SG9 |
| M98_enn314 | 61 | YNKLHEENERLKEEIGGYLDKQ | 4.7 | SG9 |
| M101_enn300 | 61 | YNKLLDENEKLKEEIGGYLDKQ | 4.7 | SG9 |
| M123_enn300.1 | 61 | YNKLLDENEKLKEEIGGYLDKQ | 4.7 | SG9 |
| M36_enn255 | 56 | ISKLYDENSKLIEERADLLDKL | 4.5 | SG7 |
| M43_enn257 | 56 | ISKLYDENSKLIEERADLLDKL | 4.5 | SG7 |
| M52_enn259 | 56 | ISKLYDENSKLIEERADLLDKL | 4.5 | SG7 |
| M53_enn252 | 56 | ISKLYDENSKLIEERADLLDKL | 4.5 | SG7 |
| M53_enn254 | 56 | ISKLYDENSKLIEERADLLDKL | 4.5 | SG7 |
| M53_enn256 | 56 | ISKLYDENSKLIEERADLLDKL | 4.5 | SG7 |

**M87 Pattern**

| Enn | Start | Sequence | Score | SG |
|---|---|---|---|---|
| M85_enn329 | 59 | YIRQLEEEREKLFDKVDQLEQQ | 4.5 | SG9 |
| M85_enn330 | 59 | YIRQLEEEREKLFDKVDQLEQQ | 4.5 | SG9 |
| M2_enn127 | 101 | NYYKLQDGIDALEKEKEDLKTT | 4.2 | SG1 |
| M8_enn95 | 110 | NYYKLQDGIDALEKEKEDLKTT | 4.2 | SG1 |
| M8_enn124 | 55 | NYYKLQDGIDALEKEKEDLKTT | 4.2 | SG1 |
| M9_enn129 | 101 | NYYKLQDGIDALEKEKEDLKTT | 4.2 | SG1 |
| M25_enn83 | 104 | NYYKLQDGIDALEKEKEDLKTT | 4.2 | SG1 |
| M25_enn82 | 104 | NYYKLQDGIDALEKEKEDLKTT | 4.2 | SG1 |

| | | | | |
|---|---|---|---|---|
| **M25_enn88** | **104** | **NYYKLQDGIDALEKEKEDLKTT** | **4.2** | **SG1** |
| **M28_enn121** | **101** | **NYYKLQDGIDALEKEKEDLKTT** | **4.2** | **SG1** |
| **M60_enn87** | **104** | **NYYKLQDGIDALEKEKEDLKTT** | **4.2** | **SG1** |
| **M66_enn129** | **101** | **NYYKLQDGIDALEKEKEDLKTT** | **4.2** | **SG1** |
| **M68_enn101** | **100** | **NYYKLQDGIDALEKEKEDLKTT** | **4.2** | **SG1** |
| **M73_enn115** | **101** | **NYYKLQDGIDALEKEKEDLKTT** | **4.2** | **SG1** |
| **M77_enn119** | **101** | **NYYKLQDGIDALEKENEDLKTT** | **4.2** | **SG1** |
| **M77_enn116** | **101** | **NYYKLQDGIDALEKENEDLKTT** | **4.2** | **SG1** |
| **M77_enn79** | **104** | **NYYKLQDGIDALEKEKEDLKTT** | **4.2** | **SG1** |
| **M77_enn81** | **104** | **NYYKLQDGIDALEKEKEDLKTT** | **4.2** | **SG1** |
| **M77_enn118** | **101** | **NYYKLQDGIDALEKENEDLKTT** | **4.2** | **SG1** |
| **M77_enn80** | **104** | **NYYKLQDGIDALEKEKEDLKTT** | **4.2** | **SG1** |
| **M79_enn114** | **103** | **NYYKLQDGIDALEKEKEDLKTT** | **4.2** | **SG1** |
| **M82_enn107** | **103** | **NYYKLQDGIDALEKEKEDLKTT** | **4.2** | **SG1** |
| **M87_enn92** | **104** | **NYYKLQDGIDALEKEKEDLKTT** | **4.2** | **SG1** |
| **M92_enn97** | **100** | **NYYKLQDGIDALEKEKEDLKTT** | **4.2** | **SG1** |
| **M92_enn100** | **100** | **NYYKLQDGIDALEKEKEDLKTT** | **4.2** | **SG1** |
| **M103_enn102** | **100** | **NYYKLQDGIDALEKEKEDLKTT** | **4.2** | **SG1** |
| **M117_enn120** | **101** | **NYYKLQDGIDALEKEKEDLKTT** | **4.2** | **SG1** |
| **M43_enn298** | **68** | **LYDDYNELQEKSAEYLERIGEL** | **4** | **SG9** |
| **M91_enn292** | **68** | **LYDDYNELQDKSAEYLERIGEL** | **4** | **SG8** |
| **M111_enn158** | **57** | **IYEELETKYEELQTKHEELLGE** | **4** | **outlier** |
| M64_enn306 | 77 | GYLDKQEQLEQLERQYQIAADK | 3.7 | SG9 |
| M80_enn311 | 77 | GYLDKQEQLEQLERQYQIAADK | 3.7 | SG9 |

**Table S4. C4BP-Binding Patterns in Strep A Strains**

| Type | M | | | | Enn | | | |
|---|---|---|---|---|---|---|---|---|
| | M2 | M22 | M68 | M87 | M2 | M22 | M68 | M87 |
| M2 | ✓ | | | | | ✓ | | ✓ |
| M3 | | ✓ | | ✓ | no Enn | | | |
| M4 | | ✓ | | | | ✓ | | |
| M4.1 | | ✓ | | | | ✓ | | |
| M8 | | ✓ | | ✓ | | | | ✓ |
| M9 | | ✓ | | ✓ | | ✓ | | ✓ |
| M11 | | ✓ | | | | | | |
| M13 | | ✓ | | | | | | |
| M14.5 | | | | | no Enn | | | |
| M15 | | ✓ | | | | ✓ | | |
| M18 | | | | | | ✓ | | |
| M22 | | ✓ | | | | ✓ | | |
| M24 | | | | | ✓ | | | |
| M25 | | ✓ | | | | | | ✓ |
| M27 | ✓ | | | | | | | |
| M28 | | ✓ | | | | ✓ | | ✓ |
| M29.2 | | | | | no Enn | | | |
| M31 | ✓ | | | ✓ | | | | |
| M32 | | | | | | | | |
| M33 | | | | | | ✓ | | |
| M34 | ✓ | ✓ | | ✓ | | | | |
| M36 | | | | | | ✓ | | |
| M38 | | | | | no genomic record | | | |
| M39 | | | | | ✓ | | | |
| M41.2 | | | | | ✓ | | | |
| M42 | | ✓ | | | | | | |
| M43 | | | | | ✓ | ✓ | | ✓ |
| M44 | | ✓ | | | | ✓ | | |
| M46 | | | | | | ✓ | | |
| M48 | | | | | | ✓ | | |
| M49 | ✓ | | | | | ✓ | | |
| M50 | | ✓ | | | | | | |
| M51 | | ✓ | | | | | | |
| M52 | | | | | | ✓ | | |
| M53 | | | | | | ✓ | | |
| M54 | | | | | | | | |
| M55 | | | ✓ | | no Enn | | | |
| M57 | | ✓ | | | | | | |
| M58 | | | | ✓ | | ✓ | | |
| M59 | | ✓ | | | | ✓ | | |
| M60 | | | | | | | | ✓ |
| M61 | | | | | no genomic record | | | |
| M62 | | | | | no genomic record | | | |
| M63 | | ✓ | | | | ✓ | | |
| M64 | | ✓ | | | | ✓ | | |

**C4BP-binding patterns in strains that:**

| | |
|---|---|
| | **Bind C4BP[a]** |
| (green) | Both M and Enn |
| (blue) | M only |
| (yellow) | Enn only |
| (peach) | Neither M nor Enn |
| (red) | **Does not bind C4BP[a]** |
| No Shading | **M, not tested for C4BP-binding** |

[a]Persson J, Beall B, Linse S, Lindahl G. Extreme sequence divergence but conserved ligand-binding specificity in Streptococcus pyogenes M protein. PLoS Pathog. 2006;2(5):e47. PubMed PMID: 16733543.

| | | | | | | | | |
|---|---|---|---|---|---|---|---|---|
| M65 | | ✓ | | | | | | |
| M66 | | | | | | | | ✓ |
| M67 | | ✓ | | | | | | |
| M68 | | | ✓ | | | ✓ | | ✓ |
| M69 | | | | | no genomic record | | | |
| M70 | | | | | | ✓ | | |
| M71 | | | | | | | | |
| M72 | | | | | ✓ | | | |
| M73 | ✓ | | | | | ✓ | | ✓ |
| M74 | | | | | ✓ | | | |
| M75 | | | | | | ✓ | | |
| M76 | | ✓ | | | | ✓ | | |
| M77 | ✓ | | | | | | | ✓ |
| M78 | | ✓ | | | | ✓ | | |
| M79 | | | | ✓ | | | | ✓ |
| M80 | | | | | | ✓ | | |
| M81 | | ✓ | | ✓ | | | | |
| M82 | | | | ✓ | | | | ✓ |
| M83 | | | | | | | | |
| M84 | ✓ | | | | | | | |
| M85 | | ✓ | | | | | | ✓ |
| M86 | | | | | | | | |
| M87 | | | | ✓ | | | | ✓ |
| M88 | | ✓ | | | | | | |
| M89 | | | | | | ✓ | | |
| M90 | | | | ✓ | | ✓ | | |
| M91 | | | | | ✓ | ✓ | | ✓ |
| M92 | | ✓ | | | | | | ✓ |
| M93 | | | | | | ✓ | | |
| M94 | | ✓ | | | | | | |
| M95 | ✓ | ✓ | | | | | | |
| M96 | | | | | no genomic record | | | |
| M97 | | | | | | ✓ | | |
| M98 | | | | | | ✓ | | |
| M99 | | | | | | | | |
| M100 | | | | | | | | |
| M101 | | | | | | ✓ | | |
| M102 | ✓ | | | | | ✓ | | |
| M103 | | ✓ | | ✓ | | | | ✓ |
| M104 | | | | | | ✓ | | |
| M105 | | | | | | | | |
| M106 | | | | | | ✓ | | |
| M108 | | | | | | ✓ | | |
| M109 | | ✓ | | | | ✓ | | |
| M110 | | | | | | ✓ | | |
| M111.1 | | ✓ | | | | | | ✓ |
| M112 | | | | ✓ | | ✓ | | |
| M113 | | ✓ | | | | | | |

| | | | | | | | | |
|---|---|---|---|---|---|---|---|---|
| M114 | ✓ | | | | | ✓ | | |
| M115 | | | | | no genomic record | | | |
| M116.2 | | | | | ✓ | | | |
| M117 | | | | | | | | ✓ |
| M118 | ✓ | | | | ✓ | ✓ | | |
| M119.2 | | | | | | | | |
| M120 | | | | | no genomic record | | | |
| M121 | | | | | | | | |
| M123 | | | | | | ✓ | | |
| M124 | | | | | | ✓ | | |
| M133 | | ✓ | | ✓ | | | | |
| M134 | | ✓ | | | | | | |
| M137 | ✓ | | | | | | | |
| M139 | | ✓ | | | | | | |
| M144 | | | | ✓ | | | | |
| M157 | | ✓ | | | | | | |
| M158 | | ✓ | | | | | | |
| M165 | | ✓ | | | | | | |
| M168 | | | | ✓ | | | | |
| M169 | ✓ | | | | | | | |
| M170 | | ✓ | | | | | | |
| M172 | | ✓ | | | | | | |
| M174 | | ✓ | | | | | | |
| M175 | ✓ | | | | | | | |
| M176 | | ✓ | | | | | | |
| M177 | | ✓ | | | | | | |
| M180 | | | | ✓ | | | | |
| M182 | | ✓ | | | | | | |
| M183 | | ✓ | | | | | | |
| M191 | | ✓ | | | | | | |
| M193 | | ✓ | | | | | | |
| M205 | | ✓ | | | | | | |
| M209 | | | | ✓ | | | | |
| M221 | | ✓ | | | | | | |
| M231 | | | | ✓ | | | | |
| M232 | ✓ | | | | | | | |
| M236 | | ✓ | | | | | | |
